# STING contributes to pulmonary hypertension by targeting the interferon and BMPR2 signaling through targeting F2RL3

**DOI:** 10.1101/2024.02.20.578386

**Authors:** Lin Deng, Chengrui Cao, Zongye Cai, Ziping Wang, Bin Leng, Zhen Chen, Fanhao Kong, Zhiyue Zhou, Jun He, Xiaowei Nie, Jin-Song Bian

## Abstract

**Rational/Objectives:** Pulmonary hypertension (PH) is an incurable disease characterized by pulmonary arterial remodeling. Endothelial injury and inflammation are the key triggers of the disease initiation. Recent findings suggest that STING (stimulator of interferon genes) activation plays a critical role in the endothelial dysfunction and interferon signaling. Here, we investigated the involvement of STING in the pathogenesis of PH.

**Methods:** PH patients and rodent PH model samples, Sugen5416/hypoxia (SuHx) PH model, and pulmonary artery endothelial cells (PAECs) were used to evaluate the hypothesis.

**Measurement and Main Results:** The cyclic GMP-AMP (cGAS)-STING signaling pathway was activated in the lung tissues from rodent PH models and PH patients, and the TNF-α induced PAECs *in vitro*. In particular, STING significantly elevated in the endothelial cell in PH disease settings. In SuHx mouse model, genetic knockout or pharmacological inhibition of STING prevented the progression of PH. Functionally, knockdown of STING reduced the proliferation and migration in PAECs. Mechanistically, STING transcriptional regulates its binding partner F2RL3 through STING-NF-κB axis, which activated the interferon signaling and repressed the BMPR2 signaling both *in vitro* and *in vivo*. Further analysis revealed that F2RL3 expression was increased in PH settings and identified negative feedback regulation of F2RL3/BMPR2 signaling. Accordingly, a positive correlation of expression levels between STING and F2RL3/ interferon-stimulated genes (ISGs) was observed *in vivo*.

**Conclusions:** Our findings suggest that STING activation in PAECs plays a critical role in the pathobiology of PH. Targeting STING may be a promising therapeutic strategy for preventing the development of PH.

## Introduction

Pulmonary hypertension (PH) is characterized by progressive remodeling of the distal pulmonary arteries leading to increased pulmonary arterial pressure and right heart failure (1). The disease initiation is triggered by PAECs injury and apoptosis, which can be mediated by pathological stressors such as inflammation and hypoxia (2–4). Mutations of BMPR2 (bone morphogenetic protein receptor 2) have been found in ∼70%-80% of heritable PAH and ∼20% of idiopathic PAH patients (5). However, with only ∼20% of individuals with BMPR2 gene mutations developing symptoms of PH (6). Thus, the “second-hit” hypothesis suggests that additional factors may be required to elicit the vascular pathology of PH (7). Inflammation is strongly evidenced as “second-hit “factor involved in the disease progression of PH and promoting BMPR2 mutation rodents susceptible to PH (8, 9). The mechanisms are remaining incomplete understanding.

The cGAS–STING signaling pathway is an innate immune inflammatory pathway that induces the release of Type I interferon and inflammatory factors by recognizing abnormally occurring DNA in the cytoplasm (10, 11). Subsequently, cGAS binds cytosolic DNA and catalyzed the synthesis of cyclin guanosine monophosphate-adenosine monophosphate (cGAMP), which acts as a secondary messenger binding and activation of STING(12). The activation of STING contributes to many inflammation-associated cardiopulmonary diseases (13, 14). The STING-associated vasculopathy with onset in infancy (SAVI) is due to the gain-of-function mutation of STING (15, 16). STING activation leads to the dysfunction of endothelial cells from the lung tissues of COVID-19 through excessive production of type Ⅰ IFN and inflammatory factors(17). However, the extent and consequences of cGAS-STING involvement in PH are still unknown.

Protease-activated receptor-4 (PAR-4), coded by F2RL3 (F2R like thrombin or trypsin receptor 3), is involved in proinflammatory reactions in ECs and is significantly increased by IL-1α or TNF-α stimulation in human coronary arteries (18, 19). F2RL3 deficiency has shown a protective role in pulmonary embolism through the inhibition of the thrombin signaling in platelets (20). Notably, previous studies have also reported an increase in F2RL3 expression in human PH patients (21, 22) and rat MCT PH model (23), whether and how F2RL3 contributes to PH is unknown.

In this study, we aimed to investigate the function role of STING in the pathogenies of PH. We found activation of cGAS-STING pathway in PH. STING inhibition prevented the development of PH and endothelial dysfunction through reducing interferon signaling and enhancing BMPR2 signaling through regulating of F2RL3 in PAECs. Our study suggests targeting STING as an effective therapeutic strategy to PH treatment.

## Methods

Detailed methods are included in the online supplement.

### Study approval

Animal Ethical information.

All the animal procedures were accredited by the Committee on Animal Research and Ethics of Southern University of Science and Technology.

#### Human Ethical information

The collection of all human samples was carried out after obtaining informed consent from the patients, and the procedures were approved by the Ethics Committee for the Use of Human Subjects of the Southern University of Science and Technology, in accordance with The Code of Ethics of the Helsinki Declaration of the World Medical Association for experiments involving human subjects. The PAH patient and matched healthy control samples were obtained from the Lung Transplant Group at the Affiliated Wuxi People’s Hospital of Nanjing Medical University in Wuxi, China. Healthy controls were sourced from donors who were deemed unsuitable for transplantation.

### STING knockout mice and Wild-type Mice

STING^-/-^ mice were a kind gift from Dr. Zhengfan Jiang of Peking University. Briefly, STING^-/-^ mice with a C57BL/6 background were produced using CRISPR-Cas9 gene-targeting technology, which introduced a 1-bp deletion at gRNA-1 and a 1-bp insertion at gRNA-2. This led to the formation of premature stop codons located between the two gRNAs. Inbred wild-type (WT) mice (C57 BL6/J) at 7 weeks old were obtained from Charles River (Guangzhou, China). The mice were housed at for a one-week acclimation and monitoring period before any experimental procedures were performed.

### Cell cultures and reagents

Human pulmonary artery endothelial cells (PAECs) were purchased from Promcell (C-12241). PAECs were culture in endothelial cell medium (ECM; Sciencell, 1001) supply with 100 U/mL penicillin, and 100 μg/mL streptomycin and 10% FBS. Cells at 4-8 passages were used for experiments. All cell culture were maintained at 37 in 95% air and 5% CO_2_.

### Sugen5416/chronic hypoxia model of PH

Pulmonary hypertension (PH) was induced in mice by exposure to chronic hypoxia using a hypobaric hypoxia chamber (10% O_2_) for three weeks with weekly subcutaneous injection of SU5416 (20mg/kg). Normoxic mice were exposed to atmospheric pressure. The temperature and relative humidity were constantly monitored, and cages were changed and cleaned every three days, with food and water accessible ad libitum. For H-151 prevention study, adult male C57BL/6J mice (8 weeks old) mice were received daily intraperitoneal injection of H-151(750nmol in 200µl PBS with 5% Tween-80) throughout the three weeks. On day 21, hemodynamic pressures and right ventricle hypertrophy (RVH) were measured, and tissues were harvest. For H-151 reversal study, adult male C57BL/6J mice (8 weeks old) were maintained in a hypoxic chamber (10% O2) for 21 days together with weekly injection of SU5416 (20mg/kg) to develop the PH phenotype. Then, intraperitoneal injection of H-151(750nmol in 200ul PBS with 5% Tween-80) daily from day 21 to day 34 under the hypoxia condition. On day 35, On day 21, hemodynamic pressures and right ventricle hypertrophy (RVH) were measured, and tissues were harvest.

### Right ventricular systolic pressure measurement

The mice were anesthetized using 2.5% isoflurane inhalation in O_2_. Subsequently, the mice were dissected to expose the right jugular vein, and a 1.4F Millar Mikro-Tip pressure catheter (Millar Instruments, Houston, TX) was inserted into the jugular vein and carefully advanced to the right ventricle. The catheter was then connected to a transducer unit, which was interfaced with a signal amplifier and recorder 3 (AD Instruments, Spechbach, Germany). The right ventricular systolic pressure (RVSP) was recorded continuously for a minimum period of 5 minutes in each animal and analyzed using Powerlab Pro Software (AD Instruments).

### Right Ventricular Hypertrophy measurement

The excised whole hearts were cleared of adjoining fatty tissue and blood vessels. After removing the right and left atria, the right ventricle (RV) was dissected from the left ventricle plus septum (LV + S) and both were dry blotted and weighed. Measurement of right ventricular hypertrophy (RVH) was determined by the ratio of RV over LV + S, and the ratio (RV/LV + S) was used in the PAH assessment methodology.

### Statistical Analysis

All qPCR and Western blot quantification results are expressed as mean and standard error of the mean (SEM). Statistical analysis was performed using Graphpad Prism 8 Software (California, USA) by unpaired student’s t-test when comparing two experimental groups. One-way ANOVA with multiple comparisons were applied in three groups analysis. Two-way ANOVA with multiple comparisons were applied when analyzing more than 2 variables. All statistics for individual experiments are showed in figure legends. Significance was determined as a P value of P < 0.05.

## Results

### cGAS-STING pathway is activated in the lung tissues of rodent PH models

qPCR analysis showed that the mRNA levels of *cGAS, STING, IRF3, IRF7, IFIT1, IFIT2, IFIT3*, and *CXCL10* were significantly upregulated in the lung tissues of various rodent PH models including SuHx and hypoxia mouse PH models (**Figure 1A and Figure S1A**) and SuHx, MCT and hypoxia rat PH models (**Figure 1B-D**). Western blotting demonstrated that the protein levels of cGAS, p-STING/STING, and p-IRF3/IRF3 were significantly increased in the lung tissues of SuHx (**Figure 1E**) and hypoxia (**Figure S1B**) PH mouse models. In consistence, the protein level of STING was significantly elevated in the lung tissues of SuHx, MCT (**Figure 1F**) and hypoxia (**Figure S1C**) PH rat models. Immunofluorescent staining revealed increased expression of STING in the endothelial cell layer of remodeled pulmonary arteries in SuHx and MCT rat PH models (**Figure 1 G**), which demonstrated that STING predominantly expressed in endothelial cells. Together, these results show that cGAS-STING pathway is activated in PH models and STING is the key dysregulated gene in endothelial cells in PH.

**Figure 1.**
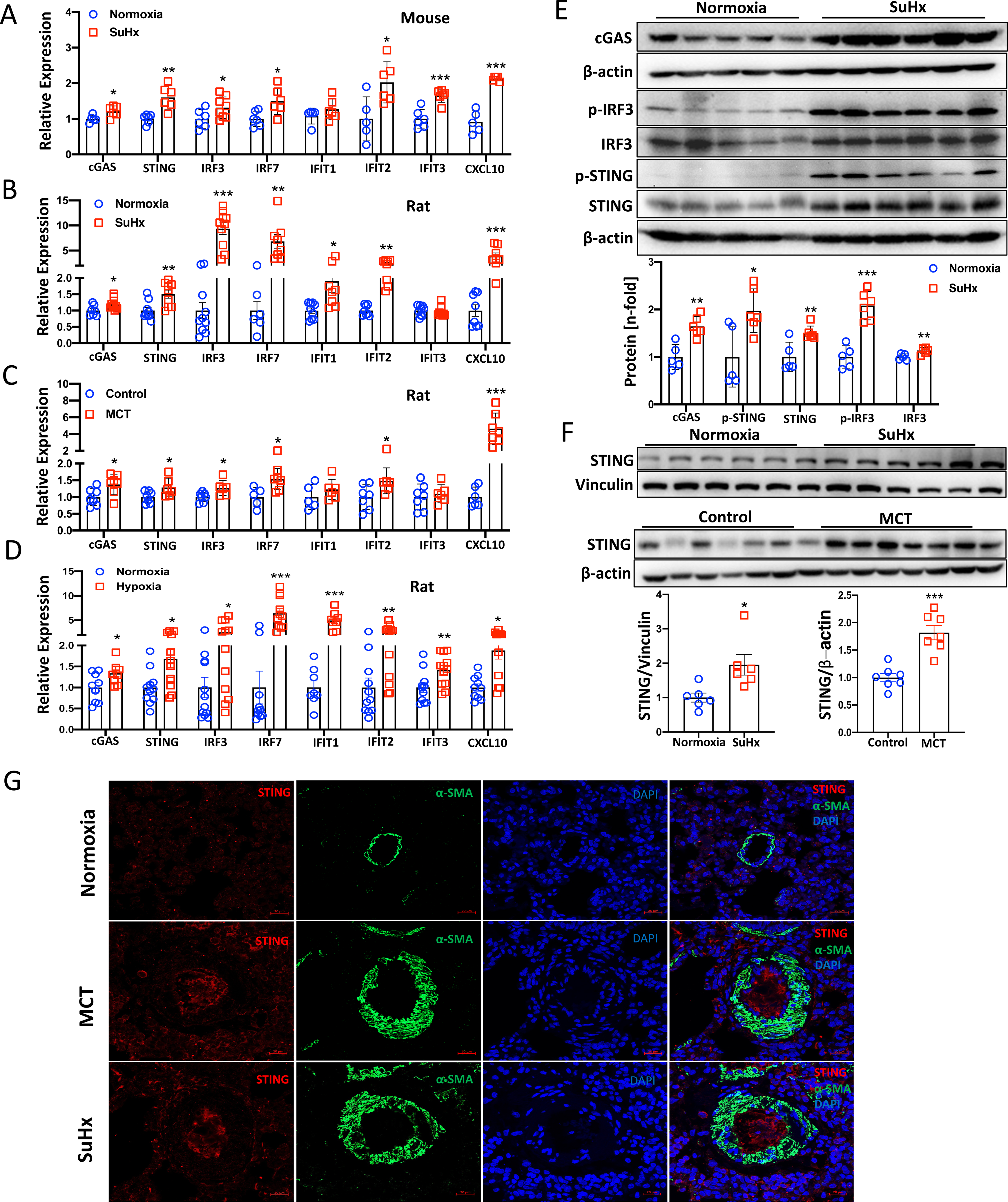
Increased cGAS-STING signaling in lung tissues of rodent PH models. **A-D**, Relative mRNA expression profiles of cGAS-STING pathway genes (*cgas, Sting, Irf3, Irf7, Ifit1, Ifit2, Ifit3* and *Cxcl10*) in SuHx mice (n=5-8) and rat (n=6-11), MCT rat (n=5-7) and hypoxic rat (n=8-12) PH models. **E**, Protein expression of cGAS-STING pathway in lung tissues of mouse SuHx PH model, and quantification of the protein expression levels of cGAS, p-STING/STING, and p-IRF3/IRF3 (n=5-6). **F**, Protein expression of STING in the lung tissues of rat MCT and SuHx PH models (n=7). **G**, Representative colocalization staining of STING (red) and α-SMA (green) in pulmonary arteries of control and MCT/SuHx rat PH models (n=4), Scale bar=20μm. Data are presented as mean ± SEM. \**P*<0.05, \*\**P*<0.01, \*\*\**P*<0.01. Statistical analysis was done using unpaired two-tailed *t* test.

### cGAS-STING pathway is activated in the lung tissues of PH patients and STING expression is increased in endothelial cells with pathologic stimulus

Further qPCR analysis showed that the mRNA levels of *cGAS, STING, IRF7, IFIT1, IFIT2, IFIT3* and *CXCL10* in the lung tissues were significantly increased in PH patients compared with those from control individuals (**Figure 2 A**). Western blotting analysis demonstrated that the expression of STING was upregulated in the lung tissues from PH patients compared with those from controls (**Figure 2 B**). Immunofluorescent and immunochemistrical staining confirmed that STING was highly expressed and increased particularly in the endothelial cell layers of the remodeled pulmonary arteries from PH patients (**Figure 2 C and Figure S1D**). This is consistent with the scRNA-Seq data from Protein Atlas (**Data not shown**) and Western blotting analysis showing that STING was highly expressed in PAECs compared with PASMCs (**Figure 2 D**). To further investigate the pathological contribution of endothelial cell STING in the development of PH, we found that *STING* expression is significantly increased in PAECs exposure to hypoxia and lung microvascular endothelial cell from hypoxia and SuHx mice PH models (**Figure 2 E-F**). Together, these findings suggest that cGAS-STING pathway is activated in PH and elevation expression of STING in endothelial cells might play critical role in the development of PH.

**Figure 2.**
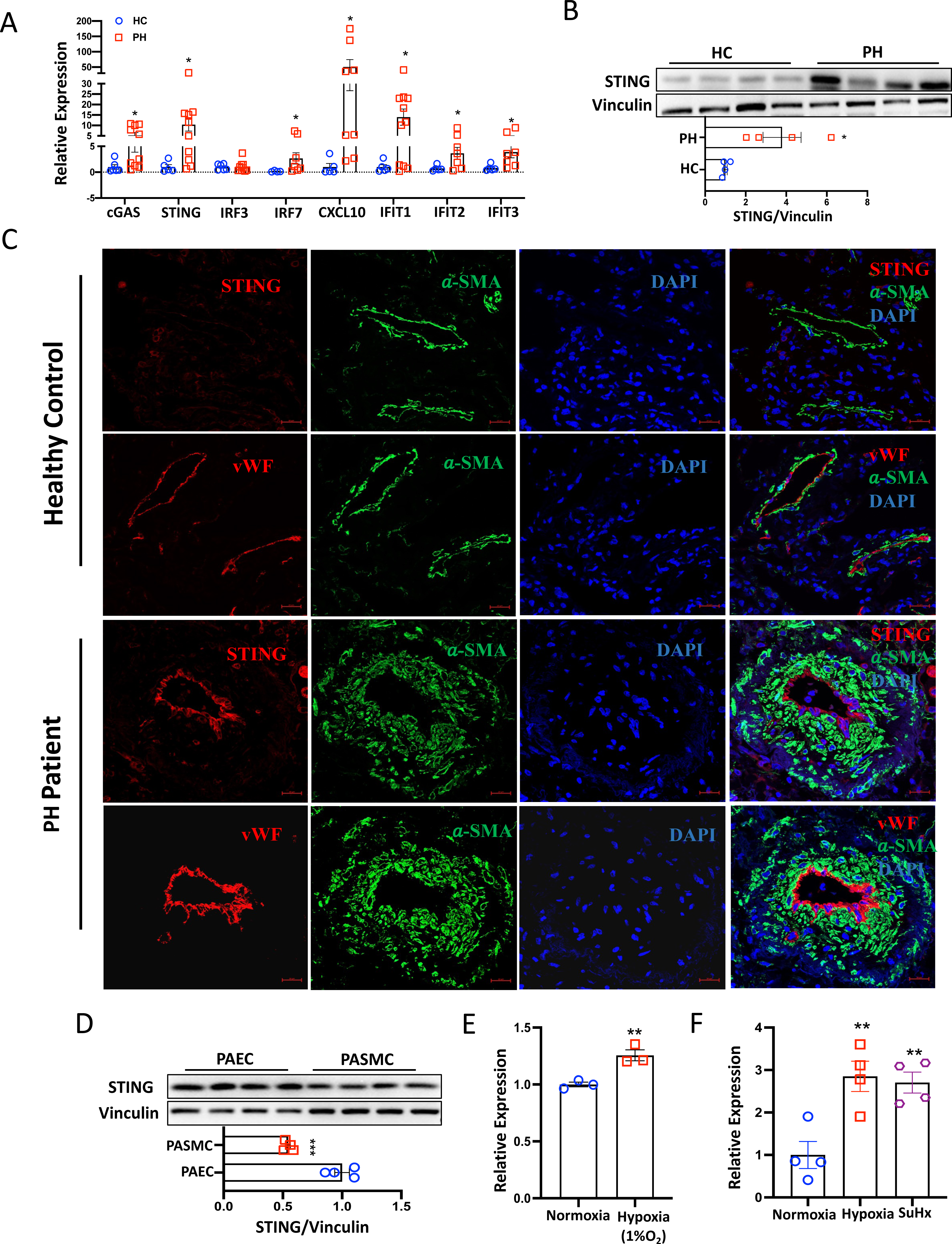
Increased cGAS-STING signaling in lung tissues of PH patients. **A**, Relative mRNA expression profiles of cGAS-STING pathway genes (*cGAS, STING, IRF3, IRF7, IFIT1, IFIT2, IFIT3* and *CXCL10*) in total lung tissues from PH patients and healthy controls (n=5-11). **B**, Protein expression levels of STING in the total lung tissues of PH patients and healthy controls (n=4). **C**, Representative colocalization staining of STING (red), vWF (red) and α-SMA (green) in pulmonary arteries of healthy controls and PH patients (n=4), Scale bar=20μm. **D**, Protein expression levels of STING in PAECs and PASMCs (n=4). **E,** Relative mRNA expression of *STING* in PAEC exposure to 1% O_2_ hypoxia for 24h (n=4). **F,** Relative mRNA expression of *Sting* in lung microvascular endothelial cells from hypoxia and SuHx mouse PH models (n=4). Data are presented as mean ± SEM. \**P*<0.05, \*\**P*<0.01, \*\*\**P*<0.01. Statistical analysis was done using Unpaired *t* test for A, B, D, and E. Statistical analysis was done using One-way ANOVA for F.

### TNF-α activates the cGAS-STING pathway and the interferon (IFN) signaling

Previous research has shown that the cGAS-STING pathway is involved in the TNF-α induced interferon responses in inflammatory arthritis (24). Here, we observed a significant increase in the transcription levels of *cGAS, STING, IRF7, IFIT1, IFIT2, IFIT3,* and *CXCL10* in PAECs with TNF-α treatment (2ng/ml) for 24 h (**Figure 3 A**). In addition, the cGAS and p-STING protein level as well as the p-IRF3 and p-TBK1 were significantly upregulated (**Figure 3 B-D**). To explore whether inhibition of STING can prevent the activation of STING dependent interferon signaling, siRNA of STING (20 μM) and H-151 (a STING inhibitor, 3μM) were applied in PAECs and followed by TNF-α treatment. It was found that STING inhibition with the two strategies significantly reduced the p-IRF3 and p-TBK1 in PAECs (**Figure 3 E-F)**. In consistence, the transcriptional levels of the interferon-stimulated genes (ISGs) were also significantly decreased with either STING silencing (**Figure 3 G**) or H-151 treatment (**Figure 3 H**). Collectively, these findings suggest that TNF-α activates the cGAS-STING pathway in PAECs, and STING inhibition prevents the activation of STING dependent interferon (IFN) signaling.

**Figure 3.**
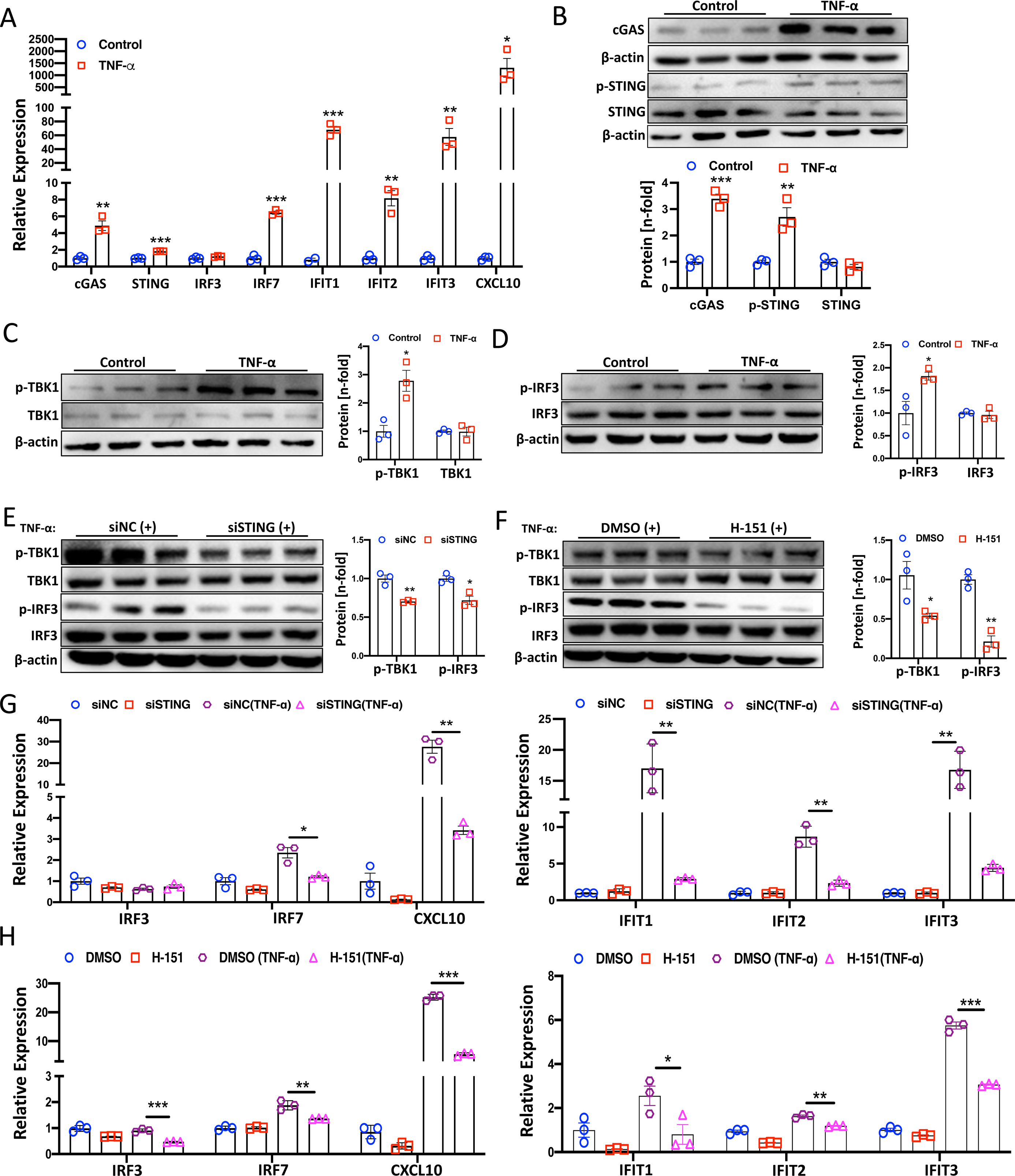
TNF-α activates cGAS-STING pathway in PAECs. **A**, Relative mRNA expression profiles of cGAS-STING pathway genes (*cGAS, STING, IRF3, IRF7, IFIT1, IFIT2, IFIT3* and *CXCL10*) in PAECs with TNF-α (2ng/ml) treatment for 24h (n=3). **B-D**, Protein expression levels of cGAS, p-STING/STING, p-IRF3/IRF3, and p-TBK1/TBK1 in PAECs with TNF-α (2ng/ml) treatment for 24 h (n=3). **E**, Protein expression levels of p-IRF3/IRF3, and p-TBK1/TBK1 in PAECs with siRNA STING and siRNA negative control (NC) transfection and followed by TNF-α (2ng/ml) treatment for 24 h (n=3). **P**, Protein expression levels of p-IRF3/IRF3, and p-TBK1/TBK1 in PAEC with H-151 (DMSO) treatment and followed by TNF-α (2ng/ml) treatment for 24 h (n=3). **G**, Relative mRNA expression profiles of interferon-stimulated genes (ISGs) (*IRF3, IRF7, IFIT1, IFIT2, IFIT3* and *CXCL10*) in PAECs transfected by siRNA STING and siRNA negative control (NC) and followed with or without TNF-α (2ng/ml) treatment for 24 h (n=3). **H**, Relative mRNA expression profiles of interferon-stimulated genes (ISGs) (*IRF3, IRF7, IFIT1, IFIT2, IFIT3* and *CXCL10*) in PAECs with H-151 (DMSO) treatment and followed with or without TNF-α (2ng/ml) treatment for 24 h (n=3). Data are presented as mean ± SEM. \**P*<0.05, \*\**P*<0.01, \*\*\**P*<0.01. Statistical analysis was done using unpaired two-tailed *t* test for A-F. Statistical analysis was done using a two-way ANOVA with multiple comparisons for G-H.

### Manipulation of STING affects the endothelial functions

EdU incorporation assays showed that STING inhibition with siRNA or H-151 significantly reduced PAEC proliferation compared to control groups (**Figure 4 A-D**). Consistently, the protein levels of proliferating cell nuclear antigen (PCNA) were also significantly decreased with STING inhibition either by siRNA or H-151 treatment (**Figure 4 E-F**). Using the scratch-would migration assay, we found that inhibition of STING decreased the cell migration rate (**Figure 4 G-H**). Additionally, we found that the expression of the endothelial cell activation marker gene ICAM-1 was significantly downregulated with siRNA STING transfection (**Figure 4 I**), indicating that endothelial activation was blocked by STING inhibition. Taken together, these *in vitro* results revealed the critical role of STING in endothelial dysfunction and activation in PAECs.

**Figure 4.**
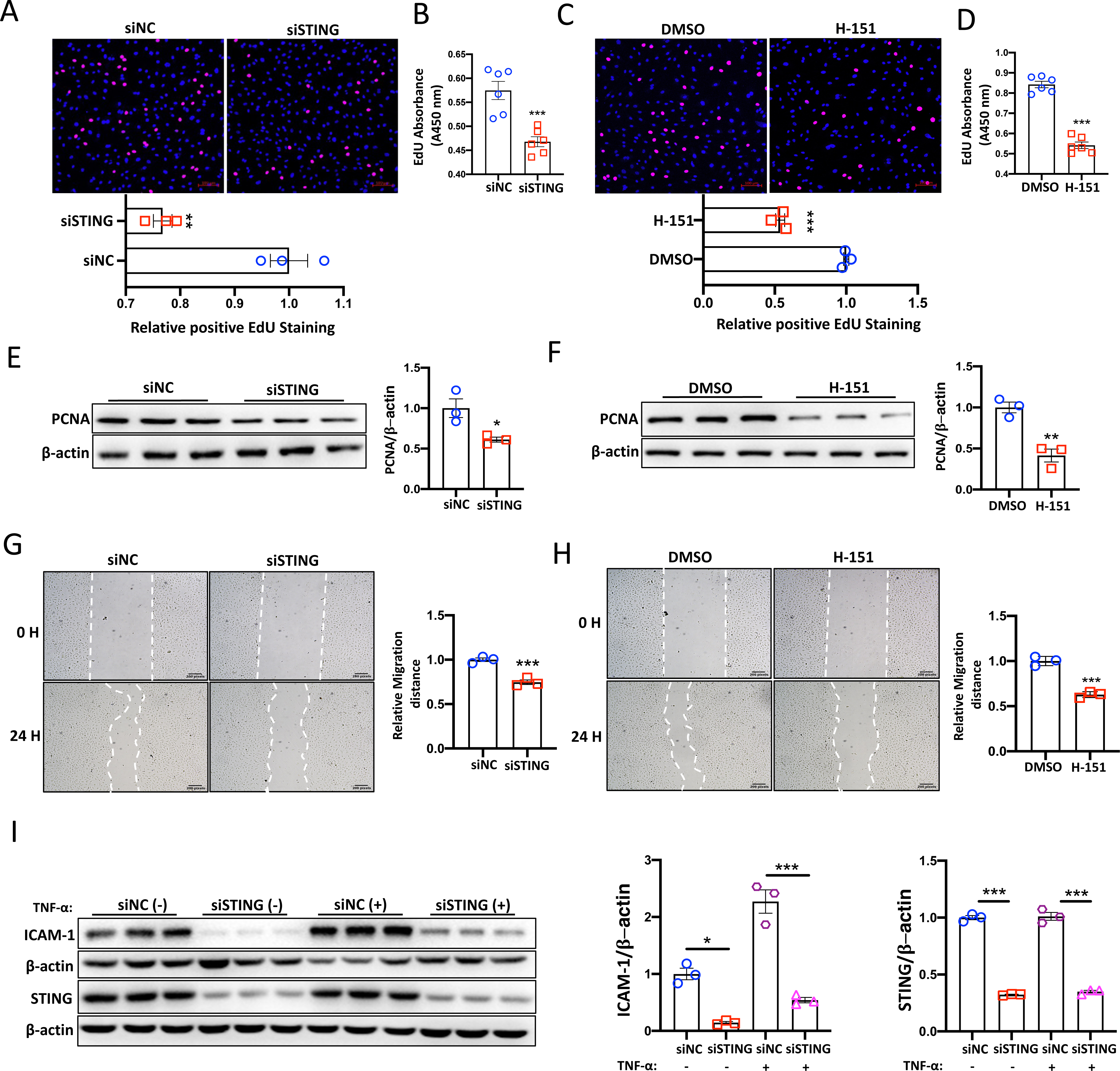
Manipulation of STING affect the endothelial function. **A** and **B**, The positive EdU staining of proliferating PAECs transfected with siRNA STING and siRNA NC were detected by fluorescent microscopy and absorbance measurement (n =3 and 6). **C** and **D**, The positive EdU staining of proliferating PAECs treated with H-151 (DMSO) were detected by fluorescent microscopy and absorbance measurement (n =3 and 6). **E** and **F**, Protein expression levels of PCNA in PAECs transfected with siRNA STING (NC) or treated with H-151(DMSO) (n=3). **G** and **H**, the relative migration distances were analyzed by wound healing assay in PAECs transfected with siRNA STING (NC) or treated with H-151(DMSO) (n=3). **I**, The protein levels ICAM-1 and STING in PAECs transfected with siRNA STING (NC) and stimulated with or without TNF-α (2ng/ml) for 24 h (n=3). Data are presented as mean ± SEM. \**P*<0.05, \*\**P*<0.01, \*\*\**P*<0.01. Statistical analysis was done using unpaired two-tailed *t* test for A-H. Statistical analysis was done using a two-way ANOVA with multiple comparisons for I.

### F2RL3 is the target of STING and transcriptional regulated by STING-NF-κB

To explore the molecular mechanisms, we performed RNA-Seq analysis on PAECs transfected with siRNA targeting STING. Totally 306 gene were significantly increased and 601 downregulated as compared control siRNA group (**Figure S2 A**). Differential expression analysis showed distinct gene expression patterns (**Figure S2 B-C**). Gene ontology analysis revealed that the silencing of STING resulted in reduced expression of genes involved in immune response, cell activation, and inflammatory response (**Figure 5 A**). KEGG pathway analysis showed several PH-related pathways were enriched, including cytokine-cytokine receptor interaction, cell adhesion molecules, complement and coagulation cascades, TNF signaling pathway, IL-17 signaling pathway, NOD-like receptor signaling pathway, and NF-kappa B signaling pathway (**Figure 5 B**). Heatmap and volcano plots revealed that genes involved in “complement and coagulation cascades” and F2RL3 gene were significantly affected (**Figure 5 C-D**). Following validation, we found that F2RL3 was significantly decreased by STING silencing (**Figure 5 E**) and this was further confirmed with Western blotting analysis (**Figure 5 F**). Previous study revealed that F2RL3 is transcriptional regulated by NF-κB p65 (25) and STING can activate downstream NF-κB signaling (26). Therefore, we propose that F2RL3 might be transcriptional regulated by STING-NF-κB axis. STING silencing significantly reduced the phosphorylation of NF-κB p65 in PAECs (**Figure 5 G**). Sequence analysis of F2RL3 2kb promoter using JASPAR (https://jaspar.elixir.no/) and NCBI (https://www.ncbi.nlm.nih.gov/gene/) identified three NF-κB potential binding sites. Chromatin immune precipitation (ChIP) assay was performed to examine the binding of NF-κB to the promoter region of F2RL3. PCR with specific primers amplifying the potential NF-κB binding sites using NF-κB ChIP samples demonstrated the binding of NF-κB in the F2RL3 promoter (**Figure 5 H**). To obtain ultimate proof for NF-κB binding to F2RL3 promoter and promoting activation, dual-luciferase reporter assay was performed. The mutation of the putative NF-κB responsive elements at site 82, 1015, 1529 of the promoter regions of F2RL3 significantly reduced the luciferase activity compared with WT (wild-type) (**Figure 5 I**), which suggest that activation of NF-κB binds to the promoter regions within the F2RL3 and increases F2RL3 transcription. Next, we separately knocked down STING and F2RL3, as well as in combination. The results indicate a significant decrease in *STING* expression upon transfection with siRNA STING or siRNA STING/F2RL3, while no changes of *STING* were observed with siRNA F2RL3 transfection in comparison to the siRNA negative control group (**Figure S3 A**). However, the *F2RL3* expression level was significantly downregulated in all three groups (**Figure S3 A**). These data suggest that F2RL3 is the downstream regulatory gene of STING. In addition, we further performed co-Immunoprecipitation (Co-IP) assay and showed the interaction between STING and F2RL3 in PAECs (**Figure 5 J**). Together, we identified F2RL3 is transcriptional regulated by STING-NF-κB axis and a binding partner of STING.

**Figure 5.**
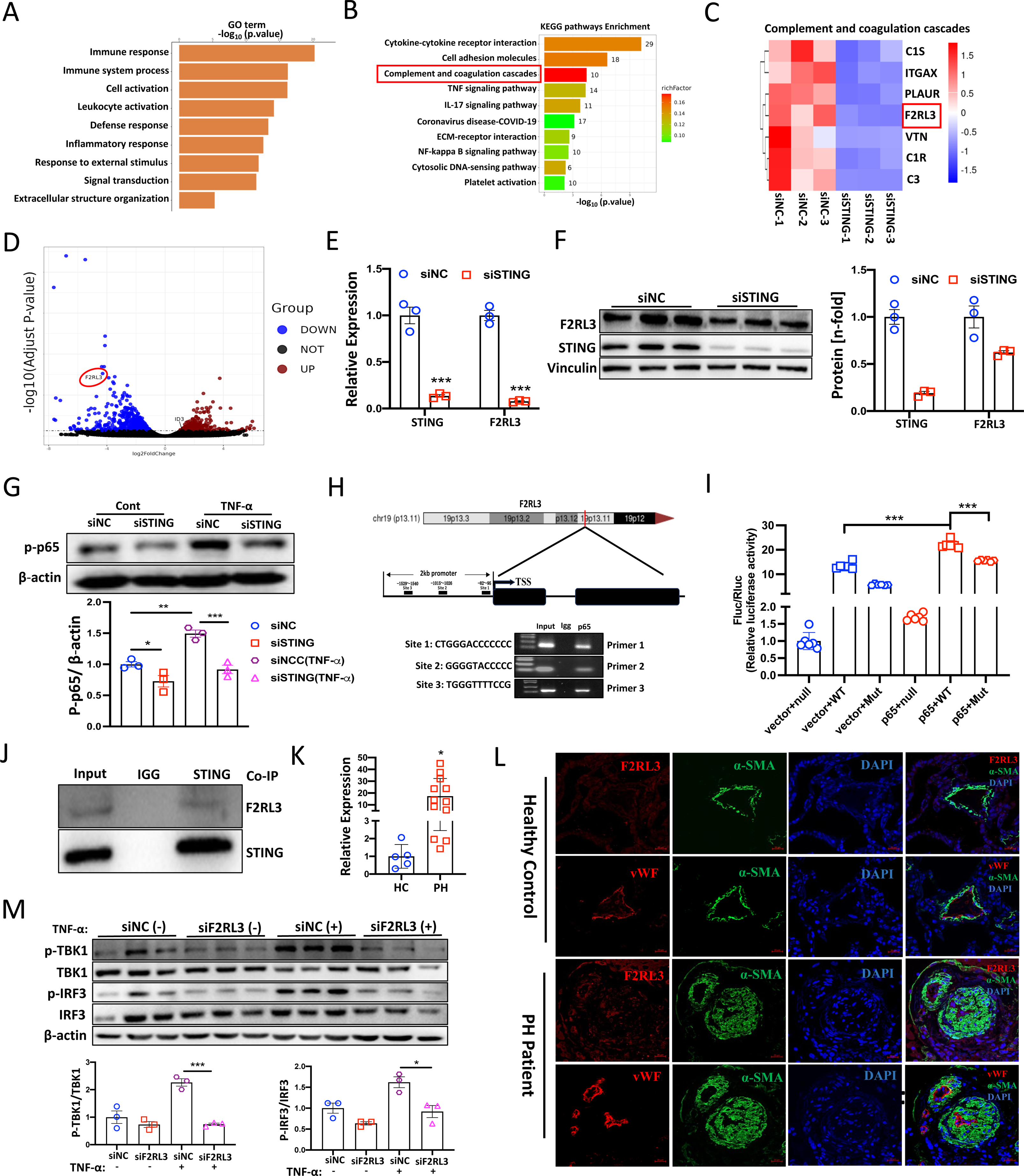
F2RL3 is transcriptional regulated by STING-NF-κB axis and involved in the pathogenesis of PH. **A** and **B**, Downregulated Gene ontology (GO) terms and KEGG pathways analysis from the bulk RNA-sequencing (RNA-seq) in PAECs with STING silencing and followed by TNF-α (2ng/ml) treatment for 24 hours. **C**, Heatmap showing dysregulated gene expression and highlighting the F2RL3 in Complement and coagulation cascades. **D**, Volcano plot showing the differentially expressed genes in PAECs and highlights the F2RL3. **E**, Relative mRNA expression of *STING* and *F2RL3* in PAECs treated with siSTING versus siNC (siNegative Control) (n=3). **F**, The protein levels of F2RL3 and STING in PAECs with siRNA STING transfection (n=3). **G,** The protein levels of phosphorylation of NF-κB p65 in PAECs treated with siSTING versus siNC±TNF-α (2ng/ml) treatment for 24 h (n=3). **H,** Schematic illustration shows the potential NF-κB binding sites of the F2RL3 promoter region, and agarose gel electrophoresis to assess the PCR reactions from the ChIP samples. **I,** Dual luciferase assay showing the transcriptional regulation of NF-κB on F2RL3 WT and mutants in 293T cells. **J,** Co-immunoprecipitation (Co-IP) in PAECs to assess the interaction between STING and F2RL3. **K,** Relative mRNA expression of *F2RL3* in lung tissues from PH patients and healthy controls (n=5-12). **L,** Representative colocalization staining of F2RL3 (red), vWF (red) and α-SMA (green) in pulmonary arteries of healthy controls and PH patients (n=4), Scale bar=20μm. **M,** The protein levels of p-STING/STING and p-TBK1/TBK1 in PAECs treated with siF2RL3 versus siNC±TNF-α (2ng/ml) treatment for 24 h (n=3). Data are presented as mean ± SEM. \**P*<0.05, \*\**P*<0.01, \*\*\**P*<0.01. Statistical analysis was done using unpaired two-tailed *t* test for E,F,and K. Statistical analysis was done using a two-way ANOVA with multiple comparisons for G,I, and M.

### F2RL3 involved in the pathogenesis of PH and TNF-α induced activation of interferon signaling pathway

To investigate the pathological relevance of F2RL3 in PH, we found that *F2RL3* was upregulated in the lung tissues from various rodent PH models (**Figure S3 B**) and PH patients (**Figure 5 K**). Immunofluorescent staining revealed that F2RL3 was particularly highly expressed in the endothelial layer of the remodeled pulmonary arteries in PH patients (**Figure 5 L)**. This finding was supported by Western blotting analysis of PASMCs and PAECs **(Figure S3 C)**, as well as the scRNA-Seq data from Protein Atlas (**Data not shown)**. Importantly, we found that *F2RL3* expression was significantly increased in PAECs exposure to hypoxia (1% O_2_) and lung microvascular endothelial cells isolated from hypoxia and SuHx mouse PH models **(Figure S3 D-E)**. Interestingly, *F2RL3* expression was significantly decreased in the lung tissues of STING^-/-^ mice compared with WT in normoxia condition *in vivo* (**Figure S3 F**) and showed strong positive correlation with the mRNA expression levels with STING (**Figure S3 G**). Mechanistically, qPCR analysis showed that both F2RL3 silencing alone and in combination with STING silencing significantly reduced the interferon-stimulated genes (ISGs) expression as indicated by *CXCL10, IFIT1, IFIT2* and *IFIT3* (**Figure S3 H)**. Furthermore, we found that F2RL3 significantly reduced the p-IRF3 and p-TBK1 in PAECs with TNF-α treatment (**Figure 5 M)**. Taken together, we found that the increasing of F2LR3 expression may contribute to the development of PH and STING-F2RL3 regulatory axis is involved in the regulating of interferon signaling.

### STING silencing enhances BMPR2 signaling by interacting with F2RL3

Apart from the downregulated pathways, STING knockdown also led to a significant induction of several PH-associated pathways, including TGF-β (GO:0071559), cellular response to TGF-β stimulus (GO: 0071560), cellular response to growth factor stimulus (GO:0071363), and positive regulation of apoptotic process (GO:0043065) signaling pathways **(Figure S4 A**). The heatmap showed genes included in the above GO pathways from RNA-Seq (**Figure S4 B**). In addition, KEGG pathway analysis revealed that the TGF-β signaling pathway was the top upregulated pathway. Most of the dysregulated genes belong to the BMP signaling (**Figure S4 A and Figure 6 A**). We therefore further performed qPCR and Western blotting experiments to validate these dysregulated BMP signaling genes. It was found that STING knockdown enhanced the mRNA expression levels of *BMPR1A, BMPR2* and *ID3* either in BMP9 (10ng/ml) (**Figure 6 B**) or TNF-α (2ng/ml) (**Figure S4 C**) stimulated conditions. Consistently, STING knockdown also led to upregulation of BMPR2 and phosphorylation of Smad1/5/9 (p-Smad1/5/9) levels in either basic condition or BMP9-(**Figure 6 C-D**)/TNF-α-(**Figure S4 D-E**) stimulated conditions. Interestingly, we also observed that silencing of F2RL3 significantly increased the p-Smad1/5/9, suggesting that F2RL3 inhibition also activates the BMP signaling (**Figure 6 E-F**). In addition, F2RL3 knockdown did not affect the protein level of STING (**Figure 6 E-F**). We further analyzed whether activation of BMP signaling affects the expression of STING or F2RL3. As shown in (**Figure 6 G**), BMP9 stimulation significantly decreased the expression of *F2RL3* rather than *STING*, whiling increasing *BMPR2, ID1* and *ID3* expression levels. Consistently, the protein levels of F2RL3 were significantly decreased with BMP9 treatment in PAECs (**Figure 6 H-I**). Furthermore, inhibition of BMPR2 significantly increased the expression of F2RL3 at both mRNA and protein levels (**Figure 6 J-L**). In order to evaluate whether STING silencing activates BMPR2 signaling is mediated by F2RL3, we performed rescue experiment by overexpression F2RL3 levels with lentivirus-F2RL3 (LV-F2RL3) infection in STING silencing conditions. Consistently, we found that STING silencing significantly increased BMPR2 and phosphorylation of p-Smad1/5/9 in STING silencing and STING silencing plus lentivirus Negative control groups. Strikingly, we observed that F2RL3 overexpression significantly blocked the elevation of BMPR2 and phosphorylation of p-Smad1/5/9 compared with the STING silencing and STING silencing plus lentivirus negative control infection (LV-NC) groups. The knockdown of STING and overexpression of F2RL3 were confirmed by Western blotting (**Figure 6 M-N**). Taken together, these data suggest that STING negatively regulates BMPR2 signaling through F2RL3, and a negative feedback loop may exist between F2RL3 and BMPR2 signaling.

**Figure 6.**
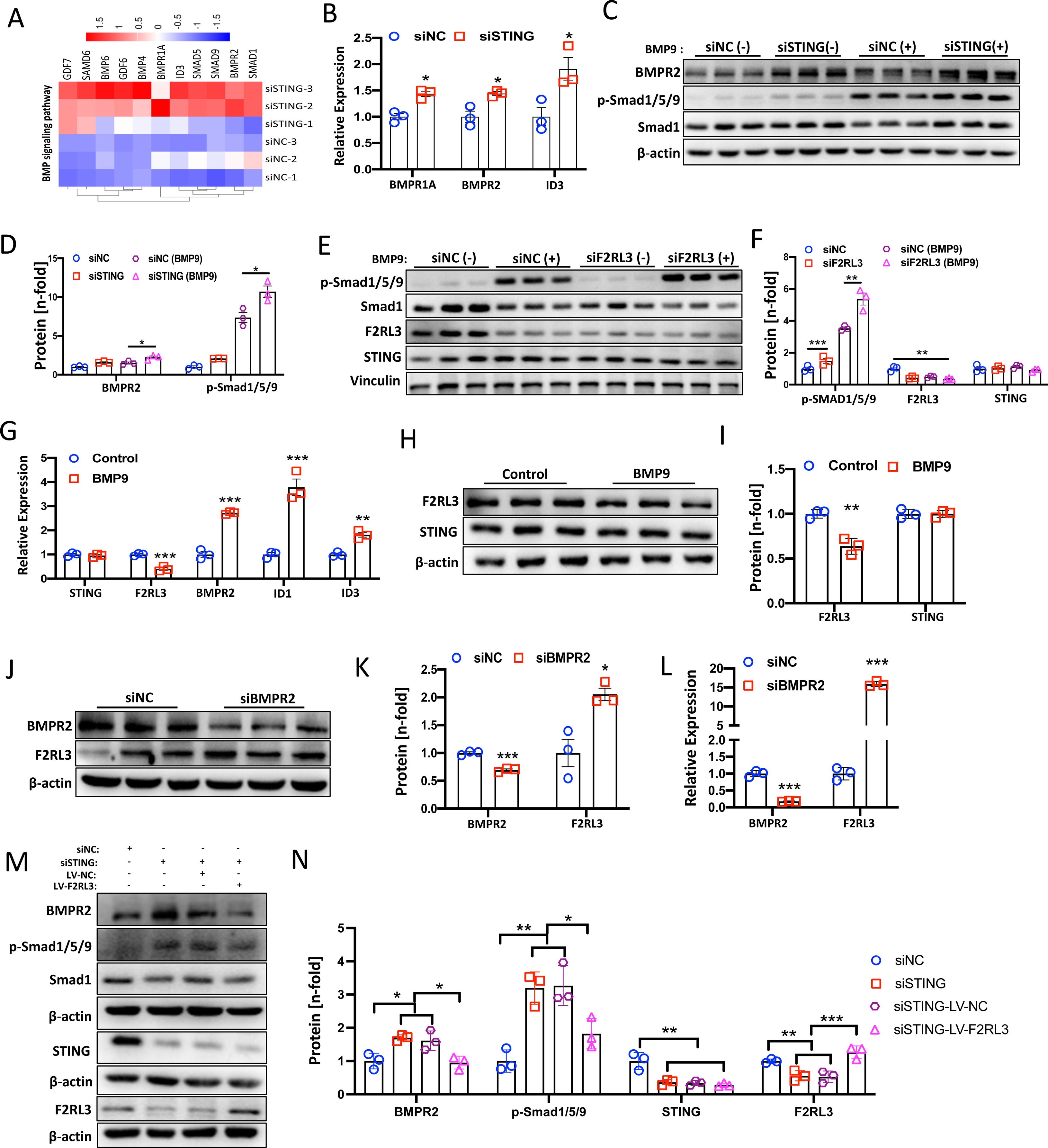
STING silencing enhanced BMPR2 signaling through interacting with F2RL3. **A**, Heatmap showing the dysregulated gene expression associated with the BMP signaling pathway. **B**, Relative mRNA expression of *BMPR1A*, *BMPR2* and *ID3* in PAECs treated with siSTING versus siNC (siNegative Control) +BMP9 (10ng/ml) treatment for 24 h (n=3). **C-D**, The protein levels of BMPR2 and p-Smad1/5/9 in PAECs treated with siSTING versus siNC±BMP9 (10ng/ml) treatment (n=3). **E-F**, The protein levels of p-Smad1/5/9, Smad1, F2RL3 and STING in PAECs treated with siF2RL3 versus siNC±BMP9 (10ng/ml) treatment for 24 h (n=3). **G**, Relative mRNA expression of *STING*, *F2RL3*, *BMPR2*, *ID1* and *ID3* in PAECs with BMP9 (10ng/ml) (n=3). **H-I**, The protein levels of F2RL3 and STING in PAEC with BMP9 (10ng/ml) treatment (n=3). **J-K**, The protein levels of BMPR2 and F2RL3 in PAECs transfected with siRNA BMPR2 and siRNA NC (n=3). **L,** Relative mRNA expression of *BMPR2* and *F2RL3* in PAECs transfected with siRNA BMPR2 and siRNA NC (n=3). **M-N**, The protein levels of BMPR2, p-Smad1/5/9, STING, and F2RL3 in PAECs transfected with siRNA STING and siRNA NC followed by lentivirus NC (LV-NC) and lentivirus F2RL3 (LV-F2RL3) infection 48 h (n=3). Data are presented as mean ± SEM. \**P*<0.05, \*\**P*<0.01, \*\*\**P*<0.01. Statistical analysis was done using unpaired two-tailed *t* test for B, G, I, K, and L. Statistical analysis was done using a two-way ANOVA with multiple comparisons for D, F, and N.

### Genetic knockout of STING alleviates the development of PH in mice

To investigate whether genetic ablation of STING *in vivo* has protective effect on the development of experimental PH. We challenged wild-type (WT) and *Sting* knockout (*Sting*^-/-^) mice with 3 weeks hypoxia together with SU5146 injection (SuHx PH model). The knockout of STING was confirmed by qPCR and Western blotting in the lung tissues (**Figure 7 A-B**), and PCR product sequencing (**Figure S5 A**). There was a significant decrease in right ventricular systolic pressure (RVSP) and right ventricular hypertrophy (RVH) in *Sting*^-/-^ mice compared with WT controls in SuHx models (**Figure 7 C-D**). No changes in heart rates were observed between groups (**Figure S5 B**). Histopathologically, *Sting*^-/-^ mice significantly reduced the pulmonary vascular remodeling compared with WT littermates (**Figure 7 E**). We found that *Sting*^-/-^ significantly improves the right ventricular (RV) function, as indicated by tricuspid annular plane systolic excursion (TAPSE) through echocardiography (**Figure 7 F**). However, there was no difference in PAT/PET ratios between two groups (**Figure 7 G**). Mechanistically, consistent with the *in vitro* data, *Sting*^-/-^ significantly decreased the p-IRF3 protein levels and enhanced the BMPR2 as well as p-Smad1/5/9 protein levels in the total lung tissues from WT and *Sting*^-/-^ mice in both normoxia and SuHx PH models (**Figure 7 H-I**). Additionally, F2RL3 protein levels were significantly reduced in Sting^-/-^ mice compared with WT mice in both normoxia and SuHx settings, while F2RL3 expression significantly elevated in WT SuHx group compared WT normoxia group (**Figure 5 S C**). Furthermore, the mRNA levels of *Sting*, *F2rl3* and ISGs expression levels were significantly increased in the total lung tissues of SuHx PH models compared with normoxia setting and reduced in the *Sting^-/-^*mice compared with WT in SuHx models (**Figure 7 J-K**). Correlation analysis revealed STING expression positively correlated with *F2rl3, Cxcl10, Irf3, Irf7*, and *Ifit2* (**Figure S5 D**). Therefore, STING knockout prevents the development of PH in mice with concomitant downregulation of F2RL3, ISGs and upregulation of BMPR2 signaling.

**Figure 7.**
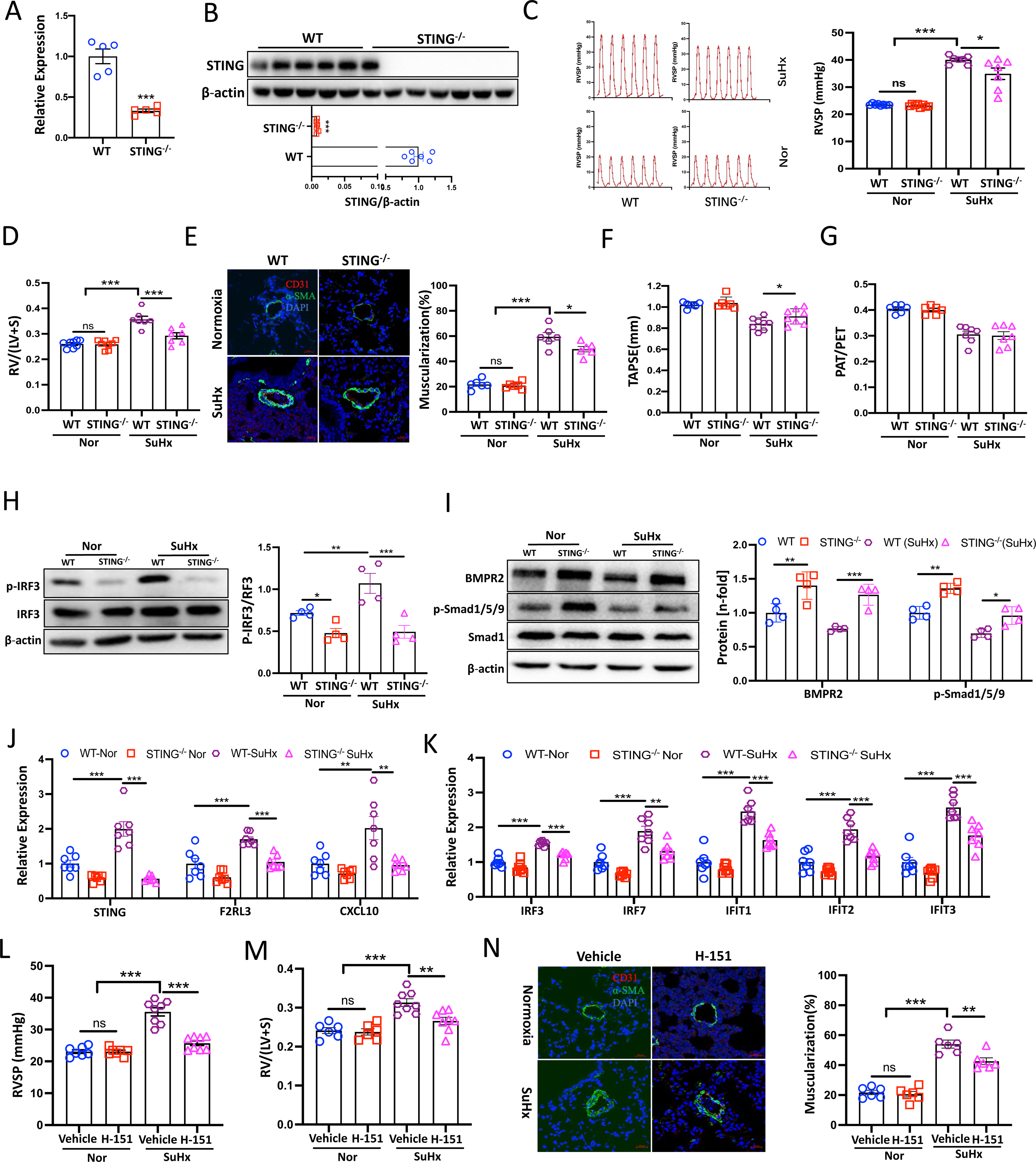
Genetic knockout and Pharmacological inhibition of STING alleviates the development of PH in mice. **A**, Relative mRNA expression of STING in the total lung tissues of WT and *Sting^-/-^* mice (n=5). **B**, The protein levels of STING in the total lung tissues of WT and *Sting^-/-^* mice (n=6-7). **C-E**, Right ventricular systolic pressure (RVSP), right ventricular hypertrophy (RVH) and pulmonary vascular remodeling in WT and *Sting^-/-^* mice in normoxia condition or exposure to hypoxia (10% O_2_) with SU5416 (20mg/kg) injection once a week for 3 consecutive weeks (SuHx) (n=6-8) Scale bar=20μm. **F-G**, Echocardiographic assessment of tricuspid annular plane systolic excursion (TAPSE) and the ratio of pulmonary artery accelerate time to ejection time (PAT/PET) in WT and *Sting^-/-^* mice under both normoxia and SuHx condition (n=6-7). **H**, The protein levels of p-IRF3 in the total lung tissues of WT and STING^-/-^ mice under normoxia and SuHx conditions (n=4). **I**, The protein levels of BMPR2 and p-Smad1/5/9 in the total lung tissues of WT and *Sting^-/-^* mice under normoxia and SuHx conditions (n=4). **J-K**, Relative mRNA expression of STING, F2RL3 and IFN-stimulated genes (ISGs) including *Irf3, Irf7, Ifit1, Ifit2, Ifit3*, and *Cxcl10* in total lung tissues of WT and *Sting^-/-^* mice under normoxia and SuHx condition (n=6). **L-N,** Right ventricular systolic pressure (RVSP), right ventricular hypertrophy (RVH) and pulmonary vascular remodeling in Vechile and H-151 treated mice in normoxia condition or exposure to hypoxia (10% O_2_) with SU5416 (20mg/kg) injection once a week for 3 consecutive weeks (SuHx) (n=6-8) Scale bar=20μm. Data are presented as mean ± SEM. \**P*<0.05, \*\**P*<0.01, \*\*\**P*<0.01. Statistical analysis was done using unpaired two-tailed *t* test for A. Statistical analysis was done using a two-way ANOVA with multiple comparisons for C-N.

### Pharmacological inhibition of STING produces preventive and therapeutic effects against PH

To examine whether pharmacological inhibition of STING can prevent PH development, WT mice received daily dose of STING covalent antagonist H-151 starting at day one exposure to chronic hypoxia together with SU5416 injection weekly (**Figure S6 A**). H-151, itself alone had no significant effect, markedly reduced the RVSP, RVH (**Figure 7 L-M**) and vascular remodeling of the distal pulmonary arteries (**Figure 7 N**) compared with vehicle group in SuHx-treated PH model. No significant difference in heart rates was found between groups (**Figure S 6 B**). Echocardiographic results demonstrated that H-151 treatment significantly improved SuHx-impaired TAPSE and PAT/PET, but itself alone had no significant effect on these two parameters under the normoxia condition (**Figure S6 C-D**). qPCR analysis revealed that H-151 treatment significantly decreased the mRNA levels of *Sting, F2rl3* and ISGs expression (**Figure S6 E**). Moreover, *Sting* positively correlated with *F2rl3, Irf3, Irf7, Ifit1, Ifit2, Ifit3,* and *Cxcl10* (**Figure S6 F-G**). Importantly, Western blotting demonstrated that p-IRF3 protein levels were significantly decreased in the H-151 SuHx group compared with vehicle SuHx group (**Figure S6 H**).

To investigate whether H-151 can be used to treat PH, mice were subcutaneously injected with SU5416 weekly and exposure to chronic hypoxia for three weeks, followed by daily injection of H-151 for another two weeks in hypoxia condition (**Figure S7 A**). Consistent with the H-151 prevention model, we found that therapeutic administration of H-151 reversed RVSP, RVH and reduced pulmonary vascular remodeling compared with vehicle controls (**Figure S7 B-D**), without affecting heart rates in both groups (**Figure S7 E**). Echocardiography showed that H-151 also improved SuHx-impaired TAPSE and PAT/PET (**Figure 8 F-G**). Mechanistically, H-151 treatment significantly decreased the mRNA expression of *Sting*, *F2rl3* and the interferon (IFN) signaling (**Figure S7 H**) and protein level of p-IRF3 (**Figure S7 I**). Moreover, STING mRNA level was positively correlated with *F2rl3, Irf3, Irf7, Ifit1, Ifit2, Ifit3*, and *Cxcl10* (**Figure S8 A**). Taken together, these results revealed that STING was a critical contributor to the pathogenesis of PH and inhibition of STING showed strong therapeutic efficacy against PH either in prophylactic or therapeutic settings.

## Discussion

In this study, we demonstrated activation of the cGAS-STING pathway in PH settings. STING played a crucial role in the development of PH with pathological features such as endothelial dysfunction, activation of interferon signaling, and repression of BMPR2 signaling both *in vitro* and *in vivo*. We also revealed that the STING transcriptional regulates its binding partner F2RL3 was involved in the activation of interferon signaling and repression of BMPR2 signaling. Additionally, the activation of BMPR2 signaling or BMPR2 silencing lead to repression or upregulation of F2RL3 in PAECs, and F2RL3 overexpression blocked the activation of BMPR2 signaling mediated by STING silencing, which suggest that silencing STING activated BMPR2 signaling through repressing F2RL3. Genetic deletion and pharmacological inhibition of STING attenuated PH development in mice. Our findings indicate that STING is a critical molecule involved in the development of PH and may serve as a potential therapeutic target.

Our data indicate that the activation of the cGAS-STING pathway in the lung tissues of various rodent PH models and PH patients. These results are consistent with the activation of cGAS-STING pathway of other cardiovascular diseases (13, 27–31). We observed the mRNA levels of *Irf3* are significantly increased in rodents PH models rather than human PH patients. The inconsistent may due to the animal PH models cannot fully mimic human disease. In addition, the PH patients we use is group 3 PH patient, which is always associated with other lung diseases. Notably, STING expression is markedly elevated in the endothelial cell layer of the remodeled pulmonary arteries and PAECs exposure to hypoxia, as well as the lung microvascular endothelial cells from hypoxia and SuHx PH models. These findings are consistent with recent studies that STING activation in lung endothelial cells plays an important role in COVID-19 (17), SAVI patients (15, 32), and acute lung injury (33). Inflammatory cytokines including IL-1β (34) and TNF-α (24) can activate cGAS-STING pathway through inducing mitochondrial DNA (mtDNA) releasing. Here, we found that TNF-α strongly activated the cGAS-STING pathway in PAECs as indicated by protein levels of p-STING, p-TBK1 and p-IRF3. Since TNF-α is a crucial driver of PH, it is possible that TNF-α mediated DNA damage or mtDNA release in PAECs might lead to the activation of the cGAS-STING pathway. Further investigation is needed to address this limitation.

Previous studies have demonstrated that IRF3 directly binds to the promoter region of ICAM-1 to induce its expression in human aortic endothelial cells (HAECs) (35), and IRF3 deficiency reduces the expression of ICAM-1 in endothelial cells *in vivo* (36). Additionally, STING gain-of-function mutation associated with the increasing expression of ICAM-1(15). Consistent with these findings, we found that TNF-α dramatically induced ICAM-1 in PAECs and STING silencing reduced the ICAM-1 expression, which suggest STING silencing prevented endothelial activation. Additionally functional data revealed that STING deficiency prevented serum-induced acceleration of PAEC proliferation and migration in classic EdU/PCNA and wound healing assays. Furthermore, STING inhibition blocked the TNF-α induced activation of interferon-stimulated genes (ISGs), such as *Ifits* and *Cxcl10* both *in vitro* and *in vivo*. These findings suggest that STING inhibition has a therapeutic effect by reducing PAECs activation, proliferation and migration, as well as interferon signaling activation. Pharmacological inhibition of STING may be a promising therapeutic strategy for reducing pulmonary vascular remodeling.

To reveal the molecular mechanisms of STING participating in the pathogenesis of PH. We performed RNA-Seq to decipher the mechanisms of STING in PAECs. We identified F2RL3 as a regulation target gene of STING. There is strong evidence linking F2RL3 DNA methylation to cardiovascular diseases such as myocardial infarction and platelet function (37). We demonstrated that STING transcriptional regulate F2RL3 though SITNG-NF-κB axis, which is consistent with previous finding (25). In addition, we confirmed that F2RL3 is a direct binding protein of STING in PAECs by Co-IP and observed a positive correlation between the expression of *Sting* and *F2rl3* in lung tissues of both wild-type and *Sting^-/-^* mice, and H-151 treated mice *in vivo*, indicating a strong correlation. We further validated the upregulation of F2RL3 expression in several rodent PH models and PH patients, as well as the lung microvascular endothelial cells from hypoxia and SuHx PH models. Like *STING, F2RL3* was found to be predominantly expressed in the endothelial cell layers in the pulmonary vasculature. Mechanistically, we observed that F2RL3 knockdown significantly reduced the levels of p-TBK1, p-IRF3, and the expression of interferon-stimulated genes (ISGs), indicating that F2RL3 is a novel player in the interferon signaling pathway. In addition, our results suggest that STING mediates the classic STING-TBK1-IRF3 signaling pathway partially through targeting F2RL3, providing a new molecular mechanism of STING action. Importantly, exploring the therapeutic potential of targeting F2RL3 *in vivo* would be valuable in the future.

PAEC injury and apoptosis mediated by pathological stressors such as inflammation and hypoxia trigger the initial stage of PH(3, 4). The dysfunction of PAEC is believed to contribute to the pulmonary vascular remodeling process either through releasing growth factors and chemokines, or activation of apoptosis-resistant pathways(38), which drive the intima and media hypertrophy by promoting the pulmonary artery smooth muscle cell (PASMC) proliferation, migration and resistant to apoptosis. We have found that STING and F2RL3 expression is highly expressed in endothelial cells. In order to further confirm the pathological relevance of STING and F2RL3, we analyzed their expression in hypoxia exposed PAECs and lung microvascular endothelial cells from hypoxia and SuHx PH models, which are classic *in vitro* and *in vivo* pathological stimuli model of PH. Both *STING* and *F2RL3* are significantly upregulated. These findings indicated that STING and F2RL3 play important pathological role of in endothelial cell of PH. However, *in vivo* PH models of STING and F2RL3 endothelial specific knockout mice should be further assessed in the future.

Recently approved ActRIIA ligand trap (Sotatercept) rebalances the activin growth differentiation factor pathway and BMP pathway to treat PH (39, 40). The enhancement of BMPR2 signaling by BMP9 shows therapeutic effect in rodents PH models, which is under pre-clinical trial (41). Therefore, enhancing BMPR2 signaling represents a promising drug target for PH treatment. In our study, we found that STING siRNA transfection in PAECs activated BMPR2 signaling. F2RL3 expression is decreased by STING silencing but not *vice versa*, which is consistent with F2RL3 is transcriptional regulated by STING-NF-κB axis. Interestingly, silencing F2RL3 activated BMPR2 signaling and BMPR2 knockdown induced *F2RL3* expression, as well as BMP9 treatment significantly repressed *F2RL3* expression rather than *STING*, all of which revealed that there is a negative feedback loop between F2RL3 and BMPR2 signaling. In addition, we demonstrated that F2RL3 overexpression repressed the STING silencing-mediated upregulation of BMPR2 signaling. Therefore, it is possible that the activation of BMPR2 signaling through STING inhibition is partially through regulating F2RL3, which revealed novel molecular mechanisms of enhancing BMPR2 signaling in PH. However, there might be other molecular mechanisms of how STING or F2RL3 regulate BMPR2 signaling in PH disease, which worth to be further explored.

Finally, in a preclinical mouse model, we demonstrated that genetic knockout of *Sting* rescued PH in the SuHx PH model, indicating the potential of targeting STING for therapy. Mechanistically, *Sting* knockout inhibited interferon signaling by reducing p-IRF3 and ISG expression, while also repressing *F2rl3* expression and inducing BMPR2 signaling. Given the interest in developing STING inhibitors for controlling innate immunity and inflammation in human diseases (42), we explored the therapeutic potential of targeting STING in PH. We found that pharmacological inhibition of STING with H-151 administration had a therapeutic effect on the development of PH in both prophylactic and therapeutic models by decreasing interferon signaling. Consistent with our findings, recently study found that pharmacological inhibition of STING with C-176 prevented the development of PH in rat SuHx model through reducing the activation of inflammasome and proinflammatory cytokines in macrophages (43). STING is ubiquitously expressed in a wide range of cells, such as macrophages, T cells and smooth muscle cells. Future research focusing on cell-specifical function and molecular mechanisms both *in vitro* and *in vivo* would be valuable for the translational application of targeting STING in the treatment of PH.

### Conclusions

We conducted a comprehensive evaluation of the cGAS-STING pathway in various rodent PH models and PH patients, revealing an important role of the STING in PH. Our findings show that STING is primarily expressed in the endothelial cells of the pulmonary vasculature and contributes to endothelial dysfunction and the pathogenesis of PH. Mechanistically, *STING* knockdown, in coordination with *F2RL3*, reduces interferon signaling and enhances BMPR2 signaling in PAECs. Translational studies using genetic deletion and pharmacological inhibition of STING show a strong therapeutic effect in preclinical mouse PH models. Together, these results highlight a novel role of *STING* in PH and suggest that it could be a promising therapeutic target for PH treatment.

## Abbreviations

|  |  |
| --- | --- |
| PH | pulmonary hypertension |
| PAH | pulmonary arterial hypertension |
| PAEC | pulmonary artery endothelial cell |
| PASMC | Pulmonary artery smooth muscle cell |
| cGAS | cyclic GMP-AMP synthase |
| STING | stimulator of interferon genes |
| BMPR2 | bone morphogenetic protein receptor type 2 |
| F2RL3 | F2R like thrombin or trypsin receptor 3 |
| RVH | right ventricle hypertrophy |
| RVSP | right ventricular systolic pressure |
| IRF3 | interferon regulatory factor 3 |
| IRF7 | interferon regulatory factor 7 |
| IFIT1 | interferon induced protein with tetratricopeptide repeats 1 |
| IFIT2 | interferon induced protein with tetratricopeptide repeats 2 |
| IFIT3 | interferon induced protein with tetratricopeptide repeats 3 |
| CXCL10 | C-X-C motif chemokine ligand 10 |
| ICAM1 | intercellular adhesion molecule 1 |
| ISG | Interferon-stimulated gene |

## Author contributions

**LD**- contributed to the concept and design of the study, data collection and analysis, and drafting the manuscript.

**CRC**- contributed to the methods development, data collection and analysis.

**ZYC**- contributed to the methods development, data collection and analysis.

**ZPW**- contributed to the methods development, data collection and analysis.

**BL**- contributed to the methods development, data collection and analysis.

**ZC**- contributed to the methods development, data collection and analysis.

**FHK**- contributed to the methods development, data collection and analysis.

**ZYZ**-contributed to the methods development, data collection and analysis.

**JH**- contributed to the methods development, data collection and analysis.

**XWN**- contributed to the concept and design of the study and review the manuscript.

**JSB**-contributed to the concept and design of the study and review the manuscript.

