## Supplementary material and methods for "STING contributes to pulmonary hypertension by targeting the interferon and BMPR2 signaling through targeting F2RL3"

### Expanded Supplementary Materials and methods

#### Echocardiography

The mice were administered anesthesia using 2.5% isoflurane inhalation in O2 and positioned on a ThermoStar temperature monitoring system pad (37 °C, RWD, China). Subsequently, the Transthoracic echocardiography findings were acquired using a Vevo1100 ultrasound system (FUJIFILM VisualSonics Inc., Canada). Tricuspid annular plane systolic excursion (TAPSE), PA acceleration time (PAT) and PA ejection time (PET) were measured, and the subsequent echocardiography data were recorded and subjected to analysis.

#### Pulmonary Vascular Remodeling analysis

In order to assess the extent of pulmonary vascular remodelling in PH models, 3 µm frontal plane lung sections were prepared and stained immunohistochemically for α-smooth muscle actin (α-SMA) and microscopically examined for non-remodelled and remodelled pulmonary arteries < 80 µm external diameter in a blinded fashion. The arteries were considered muscularised by the presence of a visible distinct thick vascular wall. In α-SMA stained pulmonary arteries, a ratio of remodelled pulmonary arteries: total counted pulmonary arteries (remodelled +non-remodelled) were calculated as the percentage of vascular remodelling. Lung sections from 6 mice for each experimental group were assessed.

#### Immunohistochemistry

Tissue sections were de-paraffinised then rehydrated by placing in two changes of fresh xylene and followed by 100%, 95%, 70%, 50% ethanol and distilled water each for 5 min. Rehydrated sections were then immersed in 10 mM citric acid buffer at pH 6 and boiled in a microwave for 4 x 5min to perform antigen retrieval. Lung sections then were then cooled to room temperature for 20 min in the citric acid buffer, followed by washing in running tap water for 10 min. Sections were then incubated with 2% BSA/PBS for 60 min to reduce non-specific background staining in a humidified chamber at room temperature. The sections were then incubated with primary antibodies at appropriate dilutions in 2% BSA/PBS and incubated at 4 °C overnight in a humidified chamber. An immunoglobulin G (IgG) negative control was used on duplicate tissue sections at the same concentration to observe any non-specific binding. After primary antibody incubation, lung sections were washed three times in tris-buffered saline (TBS) for 10 min and the secondary antibody (DAKO-Agilent, K5007) was applied on the sections and incubated for 30 min at room temperature. After incubation, sections were then washed three times in TBS for 10 min. and then visualized by Dako REAL™ DAB+ Chromagen. To quench the reaction, sections were immersed in TBS water for 5 min. Lung sections were then counterstained with haematoxylin (Cell Pathology Ltd, Newtown Powys, UK) for 1 min and washed in running tap water for 10 min. Lung sections were dehydrated by a water/alcohol gradient consisting of 70% ethanol for 1 min, 90% ethanol for 1 min, 100% ethanol for 1 min and finally twice in fresh xylene for 5 min. The lung sections were mounted with glass coverslips using DPX non-aqueous mounting medium (Merck Millipore, Darmstadt, Germany). Images were obtained using a Leica DMI6000 B microscope (Leica Microsystems, Germany).

**Immunofluorescence**

For immunofluorescent staining, the sections were prepared from formalin-fixed, paraffin-embedded samples. Deparaffinization and rehydration were carried out by washing the sections in Xylene for 5 minutes, followed by sequential washes in 100%, 95%, 70%, and 50% ethanol for 5 minutes each. Subsequently, the sections were rinsed in water for 5 minutes. To retrieve the antigens, the sections were placed in boiling sodium citrate buffer (10mM, pH 6.0) and heated in a microwave for 10 minutes. After cooling for 15 minutes, the sections were further cooled under running water for 10 minutes and then washed twice in TBS-T for 5 minutes each. Next, the sections were blocked in 2% BSA/PBS in TBS for 1 hour at room temperature, followed by overnight incubation with the primary antibodies at 4 °C.in the following dilutions of blocking buffer: vWF (A0082, Dako, 1:100), αSMA (MO851, Dako, 1:1000) and control IgGs (Mouse IgG, Invitrogen 10400C and Rabbit IgG, Abcam ab172730), at equivalent concentrations to primary antibodies. After incubation, the sections were washed three times for 5 minutes each in TBS-T. Subsequently, the sections were incubated with fluorescent-conjugated secondary antibodies (Goat Anti-Mouse, Alexa Fluor 488, Invitrogen A11001, and Goat Anti-Rabbit, Alexa Fluor 546, Invitrogen A11010) at a dilution of 1:500 for 1 hour at room temperature. Following this, the sections were washed three times for 5 minutes each in TBS-T and then mounted in ProLongTM Gold antifade reagent with DAPI (Invitrogen, P36935).

### SiRNA transfection

PAECs were transfected with siRNAs using Lipofectamine RNAiMAX transfection Reagent (Cat NO.13778150, invitrogen). Briefly, PAECs were seeded in 6-well plates overnight with 70-90% confluence at the transfection time. The following day 20μm siRNA oligomer was diluted into 500 μl Opti-MEM® per well and mixed gently. At the same time, 3 μl Lipofectamine RNAiMAX were diluted into 500 μl Opti-MEM® and mixed gently and incubated for 5 min at room temperature. After the incubation, Lipofectamine RNAiMAX / Opti-MEM® and siRNA/ Opti-MEM® were combined and mixed gently and incubated for a further 20 min at room temperature to allow transfection complexes to form. Then, 1ml transfection complexes were added to each well without culture media and incubated at 37 ℃and 5% $CO_2$ for 6-8 h and changed with fresh full culture media after. Downstream analyses were carried out after 2-3 days. STING siRNA (sc-92042) and Control siRNA-A (sc-37007) were ordered from SantaCruz. In addition, siRNA F2RL3 (targeting sequence:), siRNA BMPR2 (targeting sequence: GCAGAAATGTCCTAGTGAA), and siRNA Negative Control were purchased from RiboBio Co., Ltd. (Guangzhou, China).

### Proliferation Assay

PAECs were treated with siSTING/siNC transfection or DMSO/H-151 followed by serum starvation for 24 hours, then cells were stimulated with full medium for proliferation. Proliferation was analysed by EdU incorporation assay using EdU assay kits (C0075S and C0088S, Beyotime, Shanghai, China) according to the manufacturer's instructions. In addition, total proteins were harvested from the treated PAECs, and western blot was applied to detect proliferating cell nuclear antigen (PCNA) protein levels.

### Wound Healing Migration Assay

Migration of PAECs was analysed using the scratch wound-healing assay in 6-well plates. All cells were serum-starved for 24 h after treatment, and vertical scratches were drawn through the confluent monolayer of cells using a P1000 pipette tip. Cells were washed with PBS to remove any cell debris caused by induction of the wound and fresh full cell culture media (10% FBS) was added to the cells. For migration assay, it is recommended to use a lower percentage of serum in the growth medium to minimize the cell proliferation, however sufficient serum is required to prevent apoptosis and/or cell detachment, which is a particular concern for transfection experiments. Scratches were imaged at 0, 10, or 12 h post scratch with three images captured for each well. Migration distance analysis was performed using Image J software where a grid composed of 10 horizontal lines was placed over the photo. The distance between the edges of the scratch wound was measured along the grid lines and a relative migrated distance was expressed as a proportion of the migrated distance based on 0 h time point. Independent experiments were performed three times, with three independent wells per condition.

**RNA extraction, reverse transcription and qPCR Analysis**

PAECs Total RNA isolation was performed using TRIzol® reagent (Invitrogen) and according to the manufacturer's instructions. The lung tissues from rodents and human were lysed using 1000 µl TRIzol® reagent (Invitrogen) and disrupted and homogenised using 5 mm stainless steel beads for 1-2 min. All RNA samples were stored at -80°C until required. cDNA was synthesised from total RNA (1000 ng per reaction) using the SuperScript™ II Reverse Transcriptase (Catalog number: 18064014, invitrogen) according to the manufacturer's instructions. After synthesis, all samples were stored at -20°C until required. Target dependant, quantitative real-time polymerase chain reaction (qRT-PCR) was later performed using Power SYBR™ Green PCR Master Mix (Life Technologies, Catalog number: 4367659). SYBR Green based qRT-CR analysis primers were synthesized commercially (Shengong Co., Ltd., Shanghai, China), each designed to target unique, transcript specific, Human Ubiquitin protein C (UBC) and mouse and rat 18S was used for RT-qPCR normalisation of genes. The gene primers used in this study were in Table 1.

**Pulmonary microvascular endothelial cell isolation**

C57 normoxia mice and mice subject to hypoxia and SuHx PH models were sterilize with 70% ethanol, Open abdominal cavity and cut through diaphragm and isolate lung tissue. Place the tissue in 15ml cold isolation buffer (DMEM F12 medium with 15% FBS) in 10cm dish on ice and wash with isolation buffer to remove blood in the tissue. Then place tissue in a dry 10cm dish and mince finely with scissors on ice. Followed by incubate with collagenase A(Cat NO.10103586001,Roche) at 37 degree for 45 min, invert the tube every several minutes. Use 30ml syringe attached to a cannula to triturate the suspension 12-15 times, avoid frothing. Then pipette the suspension through a 70 μ m cell strainer into a 50ml tube, wash strainer with 10ml isolation buffer. Spin suspension at 4 degree, 400g, 8 min, followed by resuspend pellet in PBS. Use 1.5ml tubes, add 1ml suspension/per mouse/per tube, add 15µl rat anti mouse CD31(Cat NO.DIA-310, DIANOVA) coated beads (Cat NO.11035, invitrogen)to each tube and rotating for 30 min at the room temperature. Place tubes on a magnetic separator for 1 minute and then discard supernatant. Remove tubes from separator and resuspend beads in 1ml/tube isolation buffer, pipette up and down, and place tube on magnetic separator for 1min and then discard supernatant (repeat 4-5 times). Resuspend pellet into 1ml Trizol followed by RNA extraction describe before.

**Chromatin Immunoprecipitation (ChIP)**

ChIP was conducted on PAECs. Cross-linking was achieved with 1% formaldehyde for 5 minutes at 37°C, followed by quenching with 0.125 M glycine for 10 minutes at room temperature. The PAECs were then washed with cold PBS containing PMSF and lysed in SDS Lysis buffer from a ChIP Assay Kit. Subsequently, Bioruptor Plus was

employed for sonication at 4°C. The whole cell lysate was precleared using Protein A+G Agarose/Salmon Sperm DNA for 30 minutes at 4°C. After extracting the 2% input sample, the remaining sample was divided and incubated with anti-p65 (
Cat no : 10745-1-AP, Proteintech) or control IgG antibody (AC011; ABclonal; Wuhan, China) overnight. Protein A+G Agarose/Salmon Sperm DNA was added for a 2-hour incubation at 4°C. The beads were subsequently washed with various buffers, and DNA-protein complexes were eluted with elution buffer. De-crosslinking was performed by adding 0.2 M NaCl and heating for 4 hours at 65°C. Protein digestion with proteinase K followed, and DNA segments were purified using a DNA Purification Kit (D0033; Beyotime Biotechnology; Shanghai, China) for qPCR reactions. Refer to Supplementary Table 1 for the ChIP-PCR primer sequences.

**Luciferase assay**

The luciferase assay was conducted following established protocols. The pGL3-basic vector was obtained from Promega. A 2 kb fragment of the human F2RL3 promoter was PCR-amplified from Hela cells, and full mutation of three binding sites fragments were generated. 293T cell lines were seeded into 6-well plates (~1 × 10^5 cells/well) and cultured overnight at 37 °C. To assess the impact of mutations on the promoter's activity, 1200 ng of pGL3-F2RL3-WT or pGL3-F2RL3-mutant constructed vectors together with pLV3-CMV- NF-κB or pLV3-CMV-NC was transfected into 293T cells using lipofectamine 3000. Additionally, 20 ng of the p-RL-TK vector was co-transfected as an internal control. Following cell harvest, luciferase activity was measured using the Dual-Luciferase Reporter Assay System as per the manufacturer's instructions. The relative activity was normalized by the Firefly luciferase activity to Renilla luciferase activity ratio, and the fold change compared to the control pGL3 vector was calculated.

**Rodents PH models**

Rat hypoxia PH model: Eight-week-old healthy male Sprague–Dawley (SD) rats were randomly allocated into normoxia and hypoxia groups. The rats were subjected to either normoxic conditions (21% $O_2$) or hypoxic conditions (10% $O_2$) for a duration of 3 weeks.

Rat SuHx PH model: Eight-week-old healthy male Sprague–Dawley (SD) rats were randomly allocated into normoxia and SuHx groups. SuHx rats were subcutaneously injected with Sugen 5416 (20 mg/kg) and exposed to 10% oxygen for three weeks (SuHx) followed by two weeks normoxia.

Rat MCT PH model: Eight-week-old healthy male Sprague–Dawley (SD) rats were randomly allocated into vehicle and MCT groups. MCT rat were received one dose of MCT (60 mg/kg body weight) subcutaneously injected to rats.

Mouse hypoxia PH model: Eight-week-old healthy male C57 mice were randomly allocated into normoxia and hypoxia groups. The mice were subjected to either

normoxic conditions (21% $O_2$) or hypoxic conditions (10% $O_2$) for a duration of 3 weeks.
Throughout the experimental period, oxygen concentrations within the chambers were continually monitored using probes to ensure accurate and consistent exposure conditions.After hemodynamic measurement, the tissues were harvest accordingly.

**Lentiviral production and viral transduction**

The F2RL3 lentivirus vector, pLV-CMV-F2RL3-3x FLAG-CopGFP-Puro, and a vector control were cloned, and lentiviral vectors were generated through triple transient transfection of HEK293T cells. The transfection involved a packaging plasmid (pCMVΔ8.74), an envelope plasmid encoding the vesicular stomatitis virus (VSVg) (pMD.2G) and the pLNT/SFFV-MCS plasmid, using polyethylenimine (PEI) (Sigma-Aldrich) as described before (PMID: 33703914). Virus-containing medium was collected 48 hours post-transfection, purified by centrifugation, and concentrated using Millipore Ultra Centrifugal Filter Units (Fisher Scientific, Cat. # UFC901024). Lentiviral titer was assessed using the Lenti-X qRT-PCR Titration Kit (TaKaRa, Cat. # 631235). PAECs were transduced with the lentivirus for 48 hours, followed by protein extraction and western blot analysis.

**Western blot analysis**

PAECs and lung tissues protein lysates were prepared in RIPA buffer supplemented with protease and phosphatase inhibitors. Protein concentrations of the lysates were determined by BCA Protein Assay Kit (Thermofisher, 23235). Totally 15-30ug of protein were used for experiments. Proteins were resolved on 10–15% SDS-PAGE gels and electroblotted onto PVDF membrane. Membranes were blocked with 5% BSA or dried skimmed milk in TBS-T and incubated in primary antibodies in blocking buffer: anti-β-actin (Abcam ab6276), anti-Vinculin (Proteintech, 66305-1-Ig). Membranes were washed and incubated with HRP-conjugated secondary antibodies and blots developed with Western Chemiluminescent HRP Substrate (Merck-Millipore, WBULS0500) and imaged using iBright (Invitrogen). The antibodies used in this study were in Table 2.

**Co-immunoprecipitation (Co-IP) assay**

Protein lysates from PAECs were subjected to incubation with either the anti-STING antibody or control IgG at 4 °C for 1 hour with rotary agitation. Subsequently, Protein A/G PLUS-Agarose beads (Santa Cruz, sc-2003) were introduced to the mixture, followed by further incubation at 4 °C overnight with rotary agitation. After incubation, centrifugation was employed to capture the protein-antibody-bead complexes. Finally, the immune complexes were washed using ice-cold lysis buffer, boiled at 95 °C for 10 minutes in 2 × SDS-PAGE sample buffer, and subsequently analyzed through western blotting.

**Bulk RNA-seq Library Construction, sequencing and Analysis**

RNA sample preparation: RNA Samples for RNA-seq experiments were obtained in biological triplicates of PAECs. Brifely, PAECs were transfected with siRNA STING and siRNA NC followed by TNF-α (2ng/ml) treatment for 24h. The RNA samples were assessed using the A260/A280 absorbance ratio measurement method using a Nanodrop ND-2000 system (Thermo Scientific, USA). The RNA Integrity Number (RIN) was determined using an Agilent Bioanalyzer 4150 system (Agilent Technologies, CA, USA). Only samples that met the required criteria were deemed qualified for library construction.

Library preparation and Sequencing: Paired-end libraries were generated using the ABclonal mRNA-seq Lib Prep Kit (ABclonal, China), following the provided instructions. Initially, mRNA was isolated from 1 μg of total RNA using oligo (dT) magnetic beads. The isolated mRNA was then fragmented using divalent cations at elevated temperatures in ABclonal First Strand Synthesis Reaction Buffer. Next, first-strand cDNAs were synthesized using random hexamer primers and Reverse Transcriptase (RNase H), utilizing the mRNA fragments as templates. Subsequently, second-strand cDNA synthesis was performed using DNA polymerase I, RNAseH, buffer, and dNTPs. The resulting double-stranded cDNA fragments were ligated with adapters to prepare the paired-end library. Adaptor-ligated cDNA was subjected to PCR amplification. The PCR products were purified using the AMPure XP system, and the quality of the library was assessed using an Agilent Bioanalyzer 4150 system. Finally, the library preparations were sequenced on an Illumina Novaseq 6000 (or MGISEQ-T7) platform, generating 150 bp paired-end reads.

Data analysis: The data obtained from the Illumina (or BGI) platform served as the input for bioinformatics analysis. All analyses were conducted using a custom pipeline developed by Shanghai Applied Protein Technology. The primary software and associated parameters used in the analysis are outlined below. The initial processing of raw data (or raw reads) in fastq format involved the utilization of in-house perl scripts. During this step, the adapter sequences were eliminated, and reads with low quality were filtered out. Specifically, reads with a string quality value equal to or below 25 accounting for more than 60% of the total lines, as well as reads with an N (indicating undetermined base information) ratio exceeding 5%, were removed. The outcome of this process resulted in obtaining clean reads that were suitable for subsequent analysis.Next, the clean reads were aligned individually to the reference genome in orientation mode using the HISAT2 software (http://daehwankimlab.github.io/hisat2/) to obtain the mapped reads. To determine the number of reads mapped to each gene, FeatureCounts (http://subread.sourceforge.net/) was employed. Subsequently, the Fragments Per Kilobase of transcript per Million mapped reads (FPKM) for each gene was calculated by considering the gene's length and the count of reads mapped to it.

Differential expression and enrichment analysis:Differential expression analysis was conducted using the DESeq2 package (http://bioconductor.org/packages/release/bioc/html/DESeq2.html). Genes with |log2FC| > 1 and a Padj value of < 0.05 were identified as significantly differentially expressed genes (DEGs). To elucidate the functional enrichment of the DEGs and understand the distinctions in gene function among the samples, Gene Ontology (GO) and Kyoto Encyclopedia of Genes and Genomes (KEGG) enrichment analyses were performed. The clusterProfiler R software package was utilized for GO function enrichment and KEGG pathway analysis. Significance of enrichment was determined when the P-value was less than 0.05 for GO or KEGG functions.

Supplemental Table 1. List of primers used for this study

| **Gene name** | **Species** | **Forward-5'** | **Reverse-3'** |
|---|---|---|---|
| STING | Human | AGCATTACAACAACCTGCTACG | GTTGGGGTCAGCCATACTCAG |
| cGAS | Human | CACGAAGCCAAGACCTCCG | GTCGCACTTCAGTCTGAGCA |
| IRF3 | Human | AGAGGCTCGTGATGGTCAAG | AGGTCCACAGTATTCTCCAGG |
| IRF7 | Human | GCTGGACGTGACCATCATGTA | GGGCCGTATAGGAACGTGC |
| IFIT1 | Human | GCGCTGGGTATGCGATCTC | CAGCCTGCCTTAGGGGAAG |
| IFIT2 | Human | AAGCACCTCAAAGGGCAAAAC | TCGGCCCATGTGATAGTAGAC |
| IFIT3 | Human | AAAAGCCCAACAACCCAGAAT | CGTATTGGTTATCAGGACTCAGC |
| CXCL10 | Human | ACTGTACGCTGTACCTGCAT | ACACGTGGACAAAATTGGCT |
| F2RL3 | human | ACGCACTGGTGTCTGAGATG | GCAGAGTTGGGGGTGCTATT |
| UBC | Human | TTGCCTTGACATTCTCGATG | ATCGCTGTGATCGTCACTTG |
| STING | mouse | GGTCACCGCTCCAAATATGTAG | CAGTAGTCCAAGTTCGTGCGA |
| cGAS | mouse | GAGGCGCGGAAAGTCGTAA | TTGTCCGGTTCCTTCCTGGA |
| IRF3 | mouse | GAGAGCCGAACGAGGTTCAG | CTTCCAGGTTGACACGTCCG |
| IRF7 | mouse | GAGACTGGCTATTGGGGGAG | GACCGAAATGCTTCCAGGG |
| IFIT1 | mouse | CTGAGATGTCACTTCACATGGAA | GTGCATCCCCAATGGGTTCT |
| IFIT2 | mouse | AGTACAACGAGTAAGGAGTCACT | AGGCCAGTATGTTGCACATGG |
| IFIT3 | mouse | CCTACATAAAGCACCTAGATGGC | ATGTGATAGTAGATCCAGGCGT |
| CXCL10 | mouse | CCAAGTGCTGCCGTCATTTTC | GGCTCGCAGGGATGATTTCAA |
| 18S | mouse | AGGAATTGACGGAAGGGCACCA | GTGCAGCCCCGGACATCTAAG |
| F2RL3 | mouse | CCGCTGCTGTATCCTTTGGT | TCCTTGAGTTCTACTGTGGGAC |
| cGAS | rat | AGCTACCAAGGTGCTGTCAA | CCACGGTGACATCTGTATCTTTG |
| STING | rat | GGGTTTGGGGGCATCTTGAAA | AAAGGGCAGACAGCAGTCACA |
| CXCL10 | rat | CTGTCGTTCTCTGCCTCGTG | GGATCCCTTGAGTCCCACTCA |
| IRF3 | rat | TAAGGGAGATCGGCTGGCTG | TGATGGAGAGGTCCCCAAGG |
| IRF7 | rat | TCTGTGACCCTCAACACCCT | TGACTTGCAGTCACTGAGGC |
| IFIT1 | rat | CCTGGACAAGGTGGAGAATGT | GGCCATGGCTCGCATATAGT |

| | | | |
|---|---|---|---|
| IFIT2 | rat | GACACAGCAGTTGAGTGTGC | ATGATTCCTTACTGGCTGTACTCA |
| IFIT3 | rat | ATCGTCTGAGTGCCCACTTT | ACTCCTTGTTGACCTCACTCAT |
| F2RL3 | rat | GGACTGCCATTTGCTCCTCTA | GGTCGCGCCAAGTGCATATC |
| 18S | rat | TCAAGAACGAAAGTCGGAGG | GGACATCTAAGGGCATCAC |
| Chip-PCR-1 | Human | GAGAACAGTGGCTGCAGATG | GGAGGACTGGAGTGTGGGT |
| Chip-PCR-2 | Human | TCTGCTCACAGAGAAGACGG | TCCATGGCCACCAATGAGAA |
| Chip-PCR-3 | Human | CAGGGAGGGACAACACTGAG | GTACCATTGGATGGGGGCTC |

Supplemental Table 2. List of antibodies used for this study

| **Target antigen** | **Vendor or source** | **Catalog#** | **Working concentration** |
|---|---|---|---|
| STING (D2P2F) | Cell signaling Technology (CST) | 13647S | 1:1000 (WB), 1:100 (ICH) |
| Phospho-IRF-3 (Ser396) | Cell signaling Technology (CST) | 4947S | 1:1000 (WB) |
| Phospho-TBK1/NAK (Ser172) | Cell signaling Technology (CST) | 5483S | 1:1000 (WB) |
| cGAS (D1D3G) | Cell signaling Technology (CST) | 15102S | 1:1000 (WB) |
| TBK1/NAK (D1B4) | Cell signaling Technology (CST) | 3504S | 1:1000 (WB) |
| IRF-3 (D83B9) | Cell signaling Technology (CST) | 4302S | 1:1000 (WB) |
| Phospho-STING (Ser366) | Cell signaling Technology (CST) | 50907S | 1:1000 (WB) |
| cGAS (D3O8O) | Cell signaling Technology (CST) | 31659S | 1:1000 (WB) |
| Phospho-STING (Ser365) | Cell signaling Technology (CST) | 72971S | 1:1000 (WB) |
| BMPR2 | Proteintech | 19087-1-AP | 1:1000 (WB) |
| STING | Proteintech | 19851-1-AP | 1:1000 (WB), 1:100 (ICH) |
| STING | Novus Biologicals | NBP3-19785 | 1:1000 (WB) |
| Smad1 | Cell signaling Technology (CST) | 6944S | 1:1000 (WB) |
| p-Smad159 | Cell signaling Technology (CST) | 13820s | 1:500 (WB) |
| F2RL3 | Proteintech | 25306-1-AP | 1:1000 (WB),1:100 (ICH) |

| | | | |
|---|---|---|---|
| Rabbit Anti-Human vWF | Dako | A0082 | 1:100 (ICH) |
| Mouse Anti-Human αSMA/ACTA2 | Dako | MO851 | 1:1000 (ICH) |
| Mouse IgG | Invitrogen | 10400C | 1:100 (ICH) |
| Rabbit IgG | Abcam | ab172730 | 1:1000 (ICH) |
| Alexa Fluor 488 Goat Anti-Mouse | Invitrogen | A11001 | 1:500 (ICH) |
| Alexa Fluor 546 Goat Anti-Rabbit | Invitrogen | A11010 | 1:500 (ICH) |
| ICAM-1 | ICAM-1 (E3Q9N) | 67836S | 1:1000 (WB) |
| PCNA | PCNA (D3H8P) | 13110 | 1:1000 (WB) |
| Cleaved-caspase3 | Cell signaling Technology (CST) | 9664s | 1:1000 (WB) |
| Caspase3 | Cell signaling Technology (CST) | 9662s | 1:1000 (WB) |
| Vinculin | Proteintech | 26520-1-AP | 1:2000 (WB) |
| β-actin | Cell signaling Technology (CST) | 4970s | 1:1000 (WB) |
