## Supplementary figures for "STING contributes to pulmonary hypertension by targeting the interferon and BMPR2 signaling through targeting F2RL3"

### Supplementary Figure 1

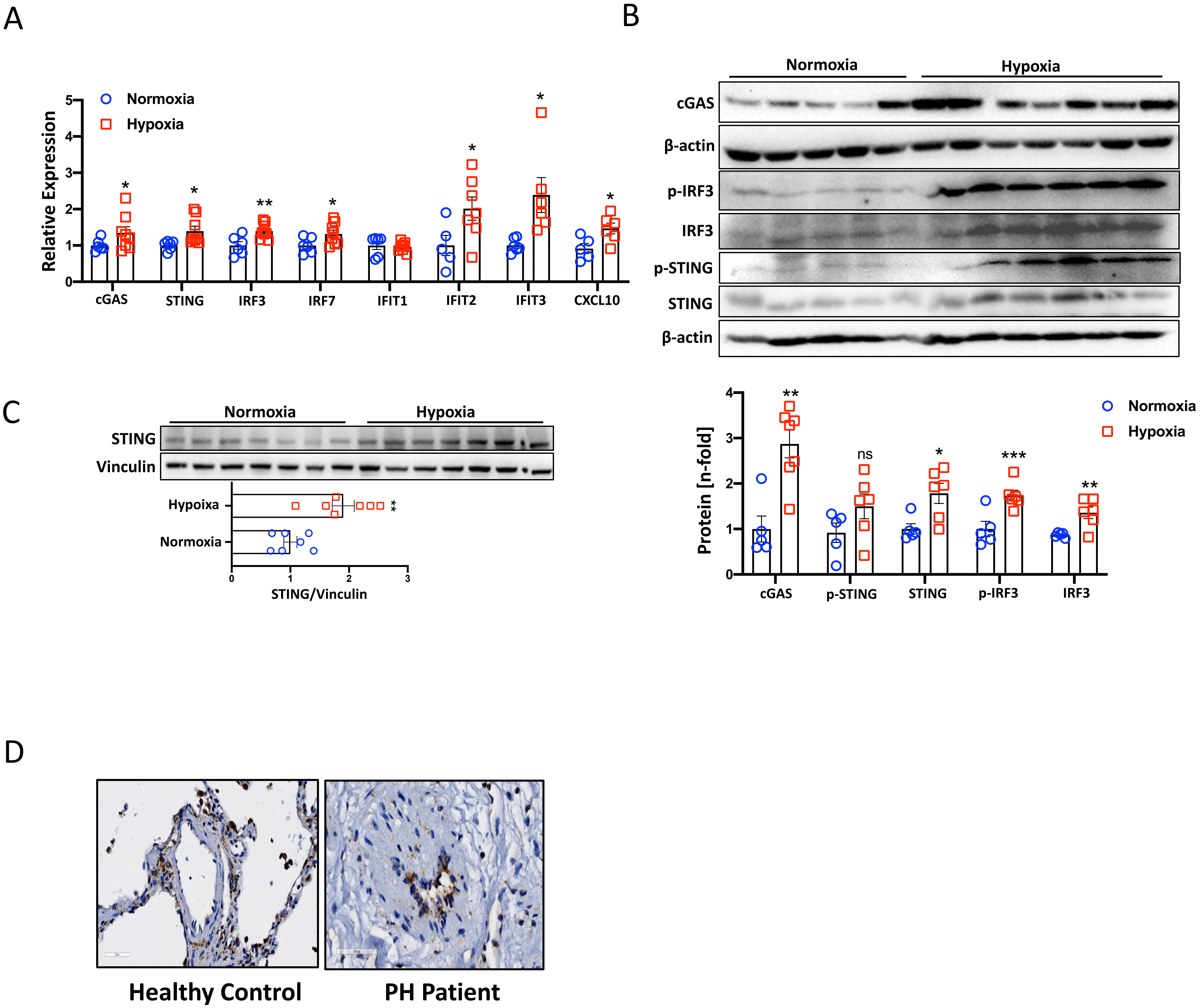

### Supplementary Figure 2

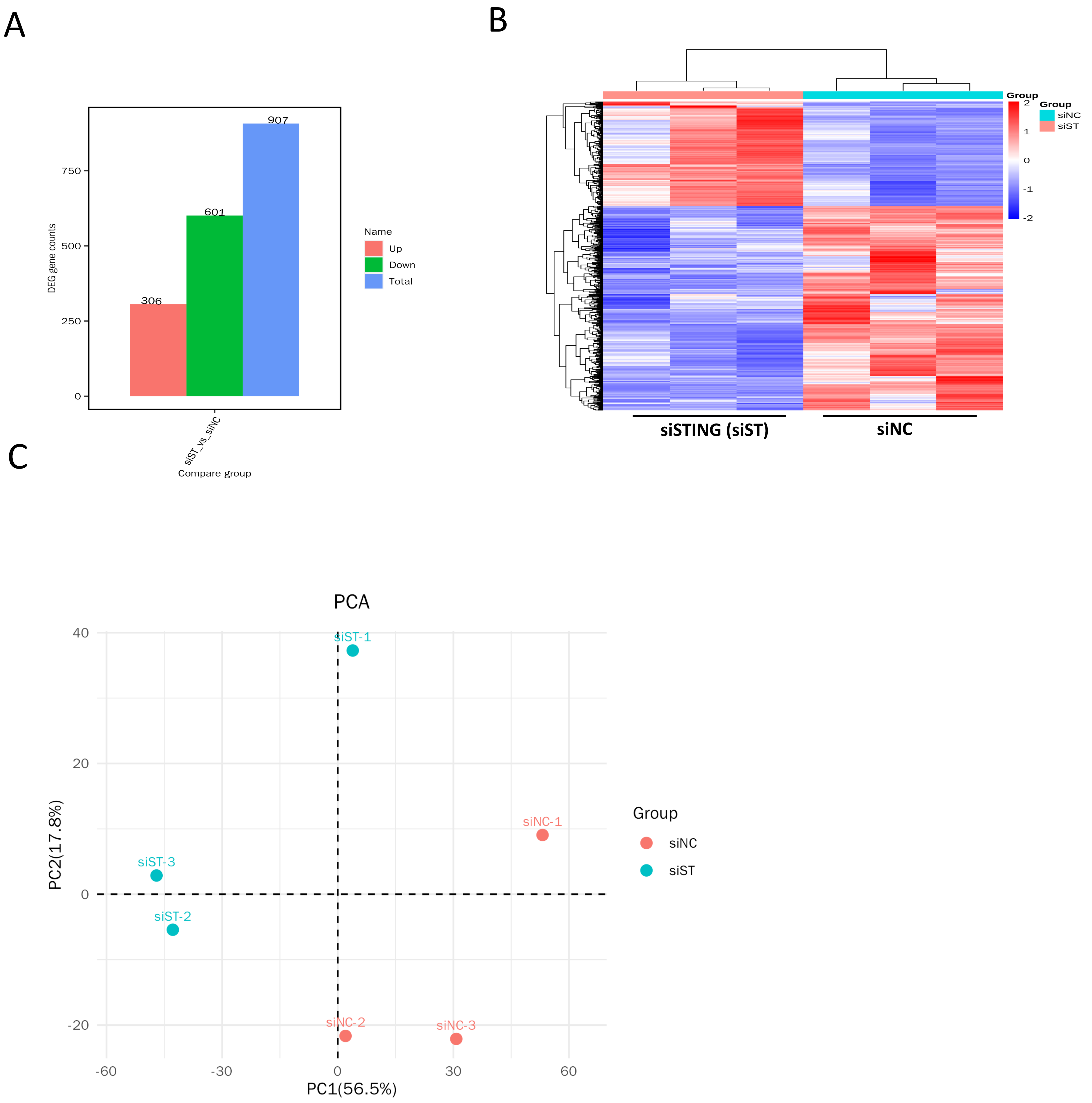

### Supplementary Figure 3

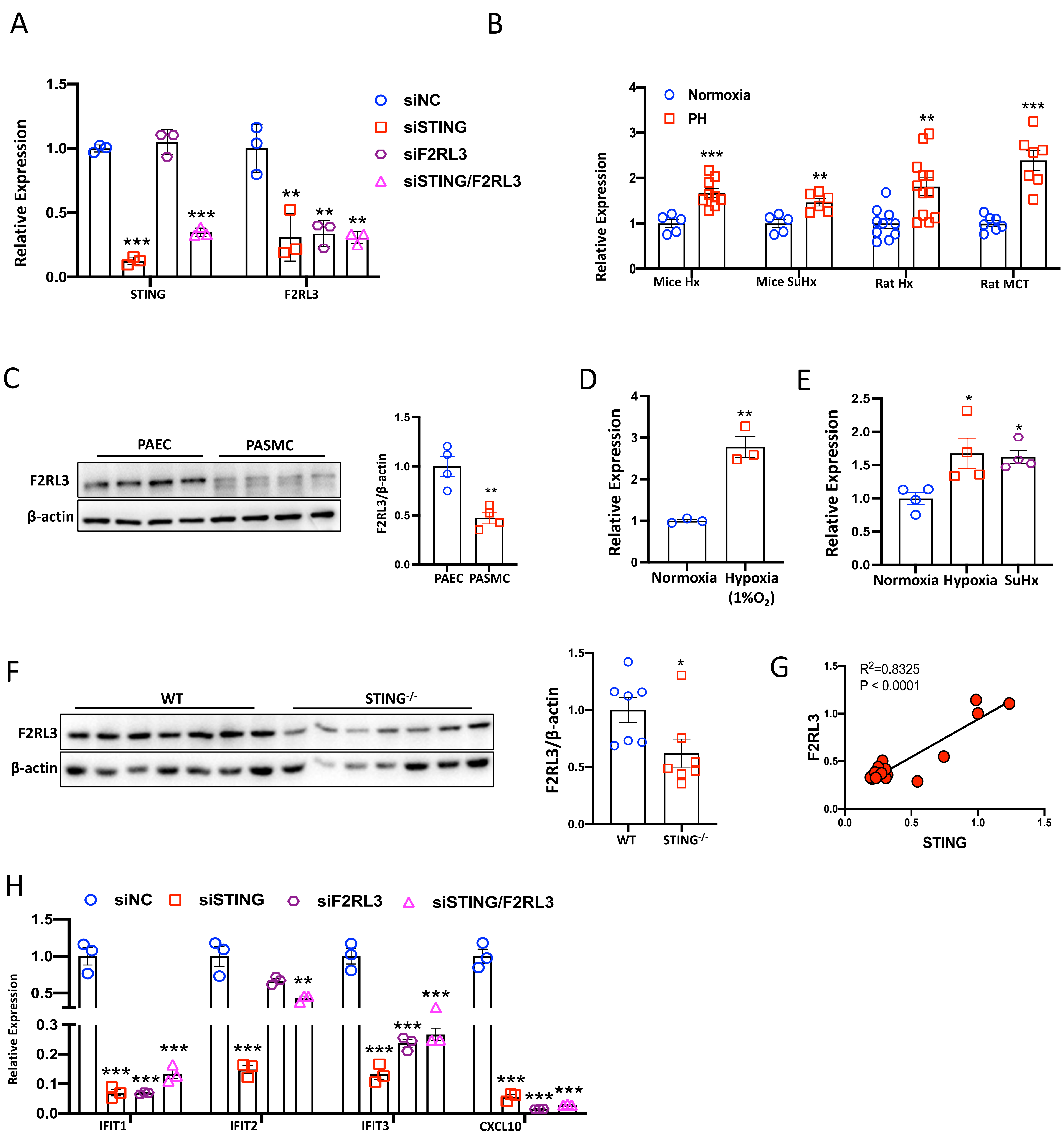

### Supplementary Figure 4

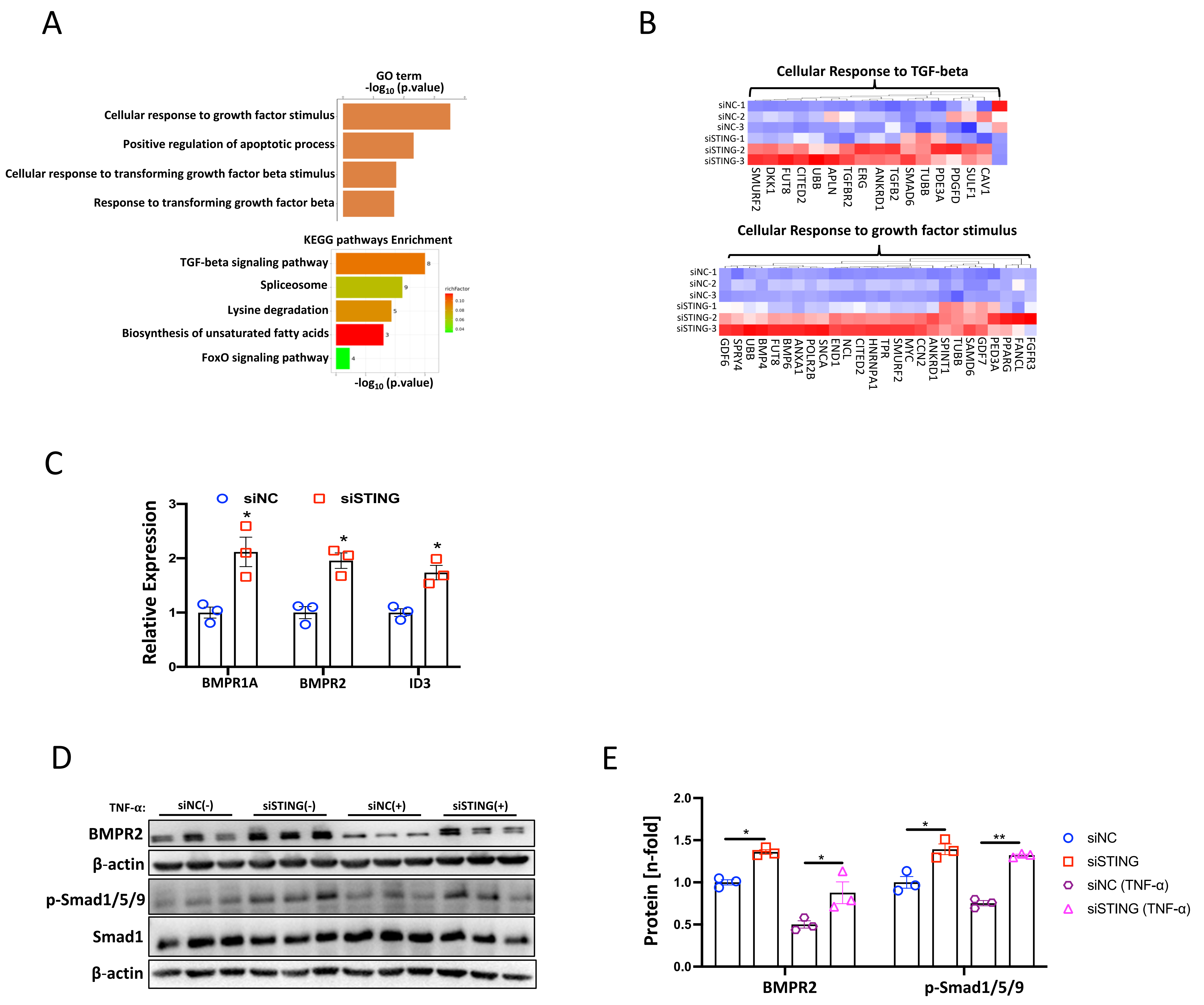

#### Supplementary Figure 5

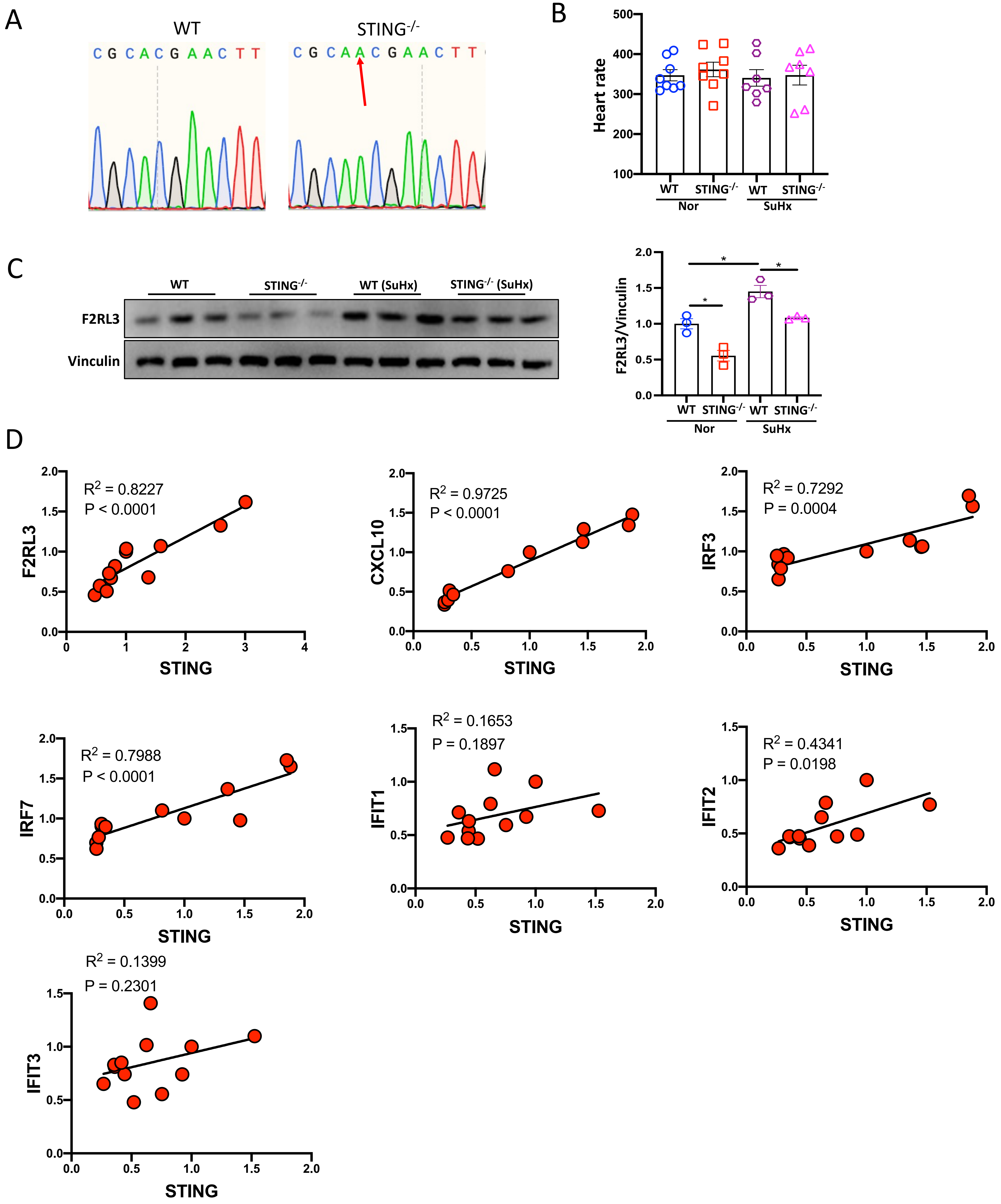

### Supplementary Figure 6

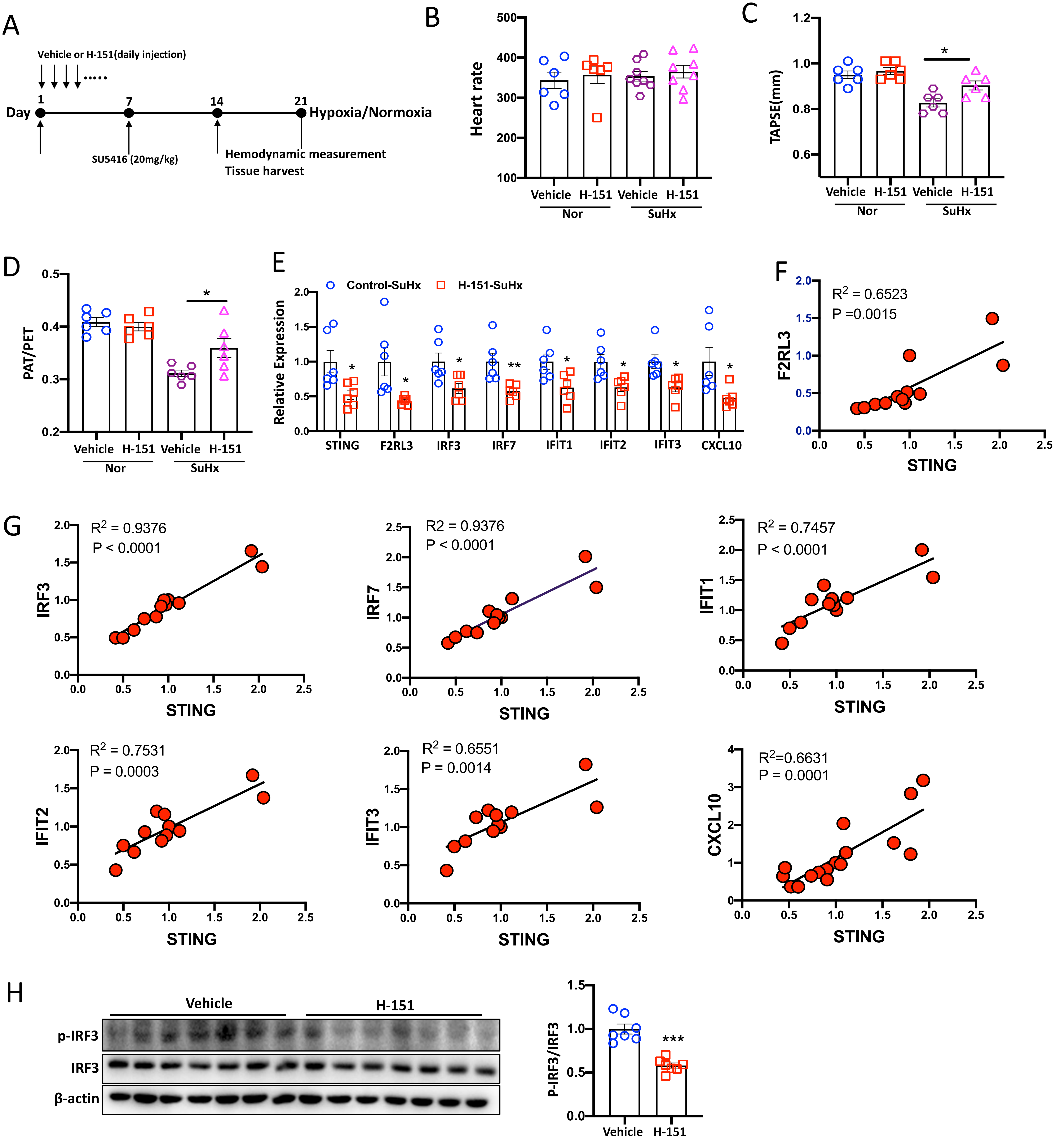

### Supplementary Figure 7

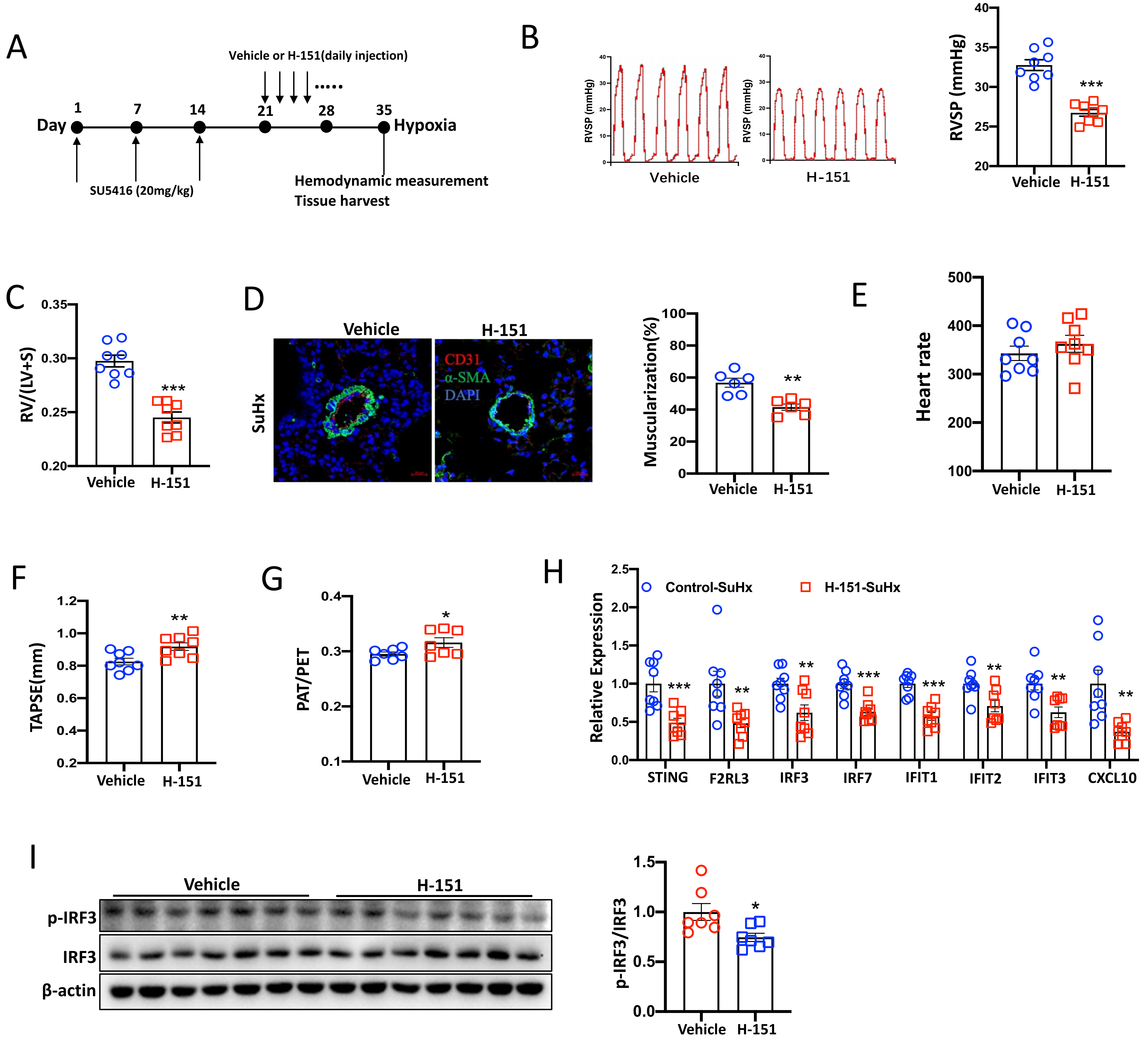

### Supplementary Figure 8

A

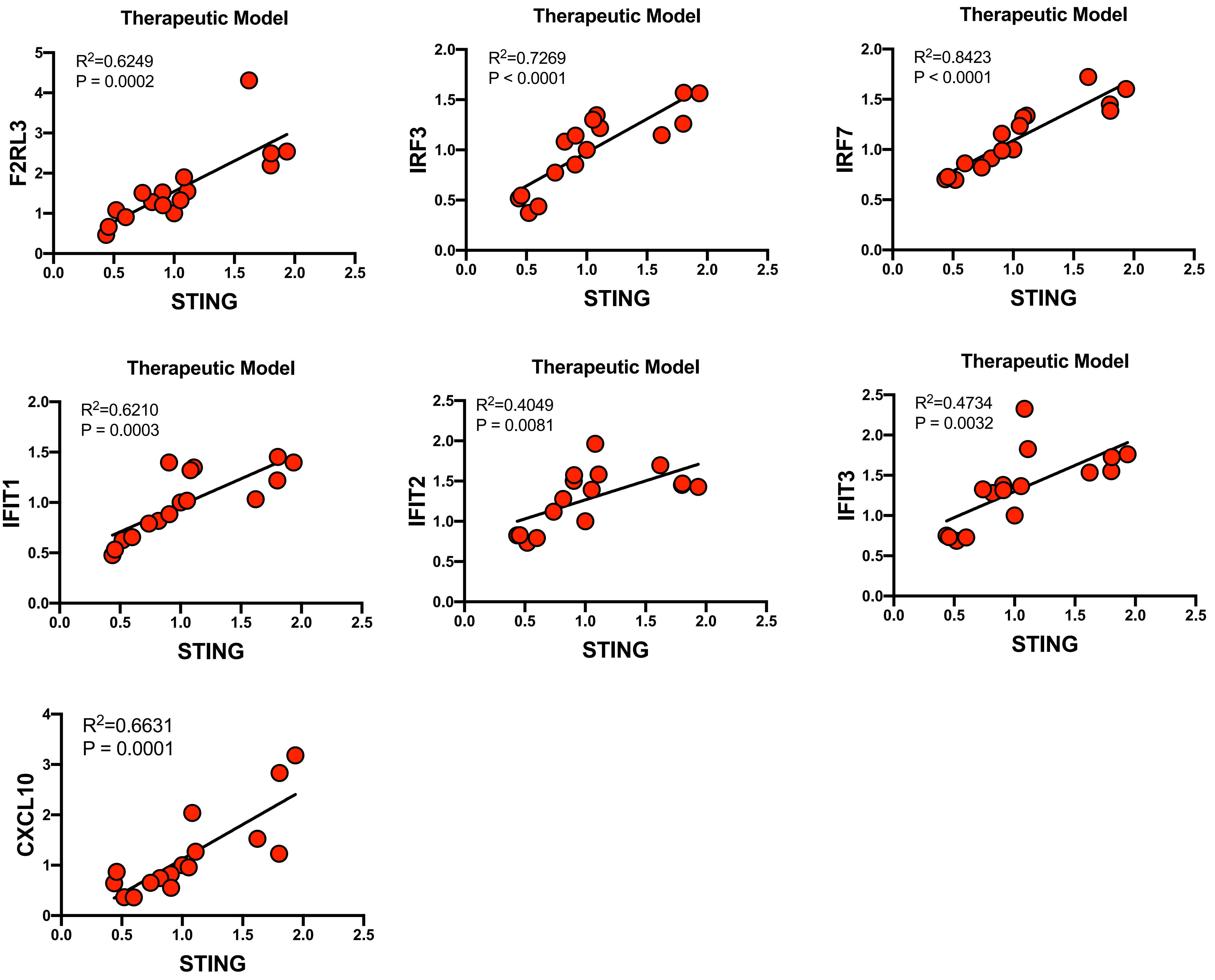

**Supplementary Figure 8**

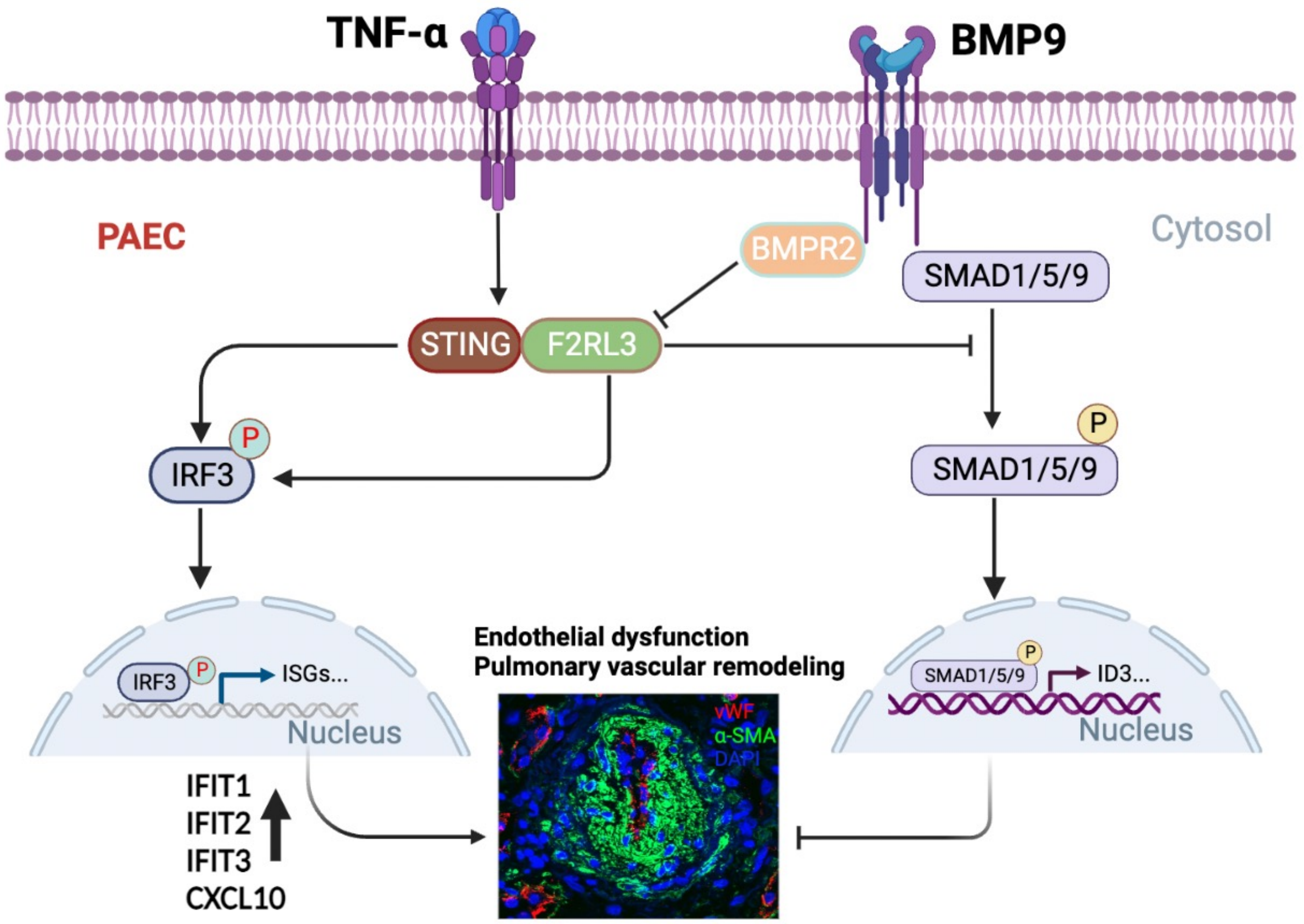
